# Immuno-Engineered Mitochondria for Efficient Therapy of Acute Organ Injuries via Modulation of Inflammation and Cell Repair

**DOI:** 10.1101/2023.06.12.544181

**Authors:** Qing Zhang, Yan Shen, Hanyi Zhang, Xuemei Li, Shengqian Yang, Chen Dai, Xiuyan Yu, Jie Lou, Chengyuan Zhang, Jinwei Feng, Chenglu Hu, Zhihua Lin, Xiaohui Li, Xing Zhou

## Abstract

Acute organ injuries represent a major public health concern, and despite recent advances in organ support therapy, managing patients with organ failure stemming from such injuries remains a formidable challenge. The pathogenesis of acute organ injuries is driven by a cascade of inflammatory reactions and mitochondrial dysfunction-mediated cell damage, two interrelated events that fuel a vicious cycle of disease progression. In this study, we engineered neutrophil membrane-fused mitochondria (nMITO) that inherit the injury-targeting and broad-spectrum anti-inflammatory activities from neutrophil membrane proteins while retaining the cell-repairing activity of mitochondria. We demonstrated that nMITO can effectively block the inflammatory cascade and replenish mitochondrial function to simultaneously modulate these two key mechanisms in diverse acute organ injuries. Furthermore, by virtue of the β-integrin inherited from neutrophils, nMITO exhibit selective homing to injured endothelial cells and can be efficiently delivered to damaged tissue cells via tunneling nanotubes, amplifying their regulatory effects on local inflammation and cell injury. In mouse models of acute myocardial injury, acute liver injury, and acute pancreatitis, nMITO effectively ameliorated immune dysfunction and repaired damaged tissues. Our findings suggest that nMITO represents a promising therapeutic strategy for managing acute organ injuries.

## Introduction

Acute organ injuries (AOIs) such as acute myocardial injury^1^, acute lung injury^2^, acute liver injury^3^, acute kidney injury^4^, acute pancreatitis^5^, and others are among the leading causes of death worldwide^6^. The rapid progression of these injuries is often driven by mitochondrial dysfunction and a highly pro-inflammatory environment, leading to organ failure, multiple organ failure, and death^7, 8^. Due to the complexity of the pathological mechanism behind AOIs, there are currently no effective clinical treatment strategies for AOI therapy. The clinical treatment of AOI mainly adopts supportive therapy for conservative treatment, which urgently requires innovative strategies with breakthrough therapeutic effects.

Mitochondrial transplantation, a novel cell repair strategy, is expected as a rising star in potentially treating of AOIs, and has shown a good cell repair ability in the animal models of AOIs including acute myocardial ischemia/reperfusion, lung injury, liver injury, stroke, and others^9, 10^. Importantly, the mitochondrial transplantation has also shown certain efficacy in clinical trials in the fields of acute myocardial ischemia/reperfusion and stroke^10^. However, the further development of mitochondrial transplantation is strictly limited by the deficiency of lesion targeting distribution and inflammation regulation ability. Mitochondrial administration in clinical research is limited to local delivery due to its inability to target lesions, greatly reducing convenience, safety, and the possibility of multiple administrations.^10^. In addition, a highly pro-inflammatory environment is also key factor driving the process of various AOI^11, 12^, while natural mitochondria can reduce inflammation to a certain extent through cell repair, they lack the ability to systemically regulate inflammation, which is insufficient for controlling the formed inflammatory cascade in AOIs.

Recently, cell-membrane-coated nanoparticles have emerged as a promising targeting delivery and therapeutic platform, which can acquire the main membrane proteins from the source cell for targeting specific lesions^13, 14^, absorbing and neutralizing complex pathological molecules^15, 16^. The advancement of cell-mimicking nanoparticles in drug delivery and detoxification inspired the development of immuno-engineered mitochondria to address the aforementioned challenges of mitochondrial transplantation. Neutrophils, a type of polymorphonuclear leukocyte, are well recognized as one of the major participants during inflammation and tissue injury^17, 18^. As neutrophils are well involved in the inflammatory cascade of AOIs^12, 19^, we herein developed a neutrophil membrane fused mitochondria (nMITO) with organ injury targeting ability, broad-spectrum anti-inflammatory activity and cell repairing effect (Fig.1a). nMITO developed in this study inherited the β integrin from neutrophil membrane on the surface, enabling efficient targeting to the injuring organ and rapid delivery to injured cells for repairing damaged mitochondrial function and injured cells. nMITO also inherited chemokine receptors and inflammatory cytokine receptors from neutrophil membrane on the surface, allowing for the absorption and neutralization of inflammatory chemokines and cytokines to reduce the local infiltration and systemic diffusion of inflammation. Through promoting cell repair and modulating inflammation, nMITO can ameliorate a variety of AOIs, including acute myocardial injury, acute liver injury, and acute pancreatitis.

**Figure 1.**
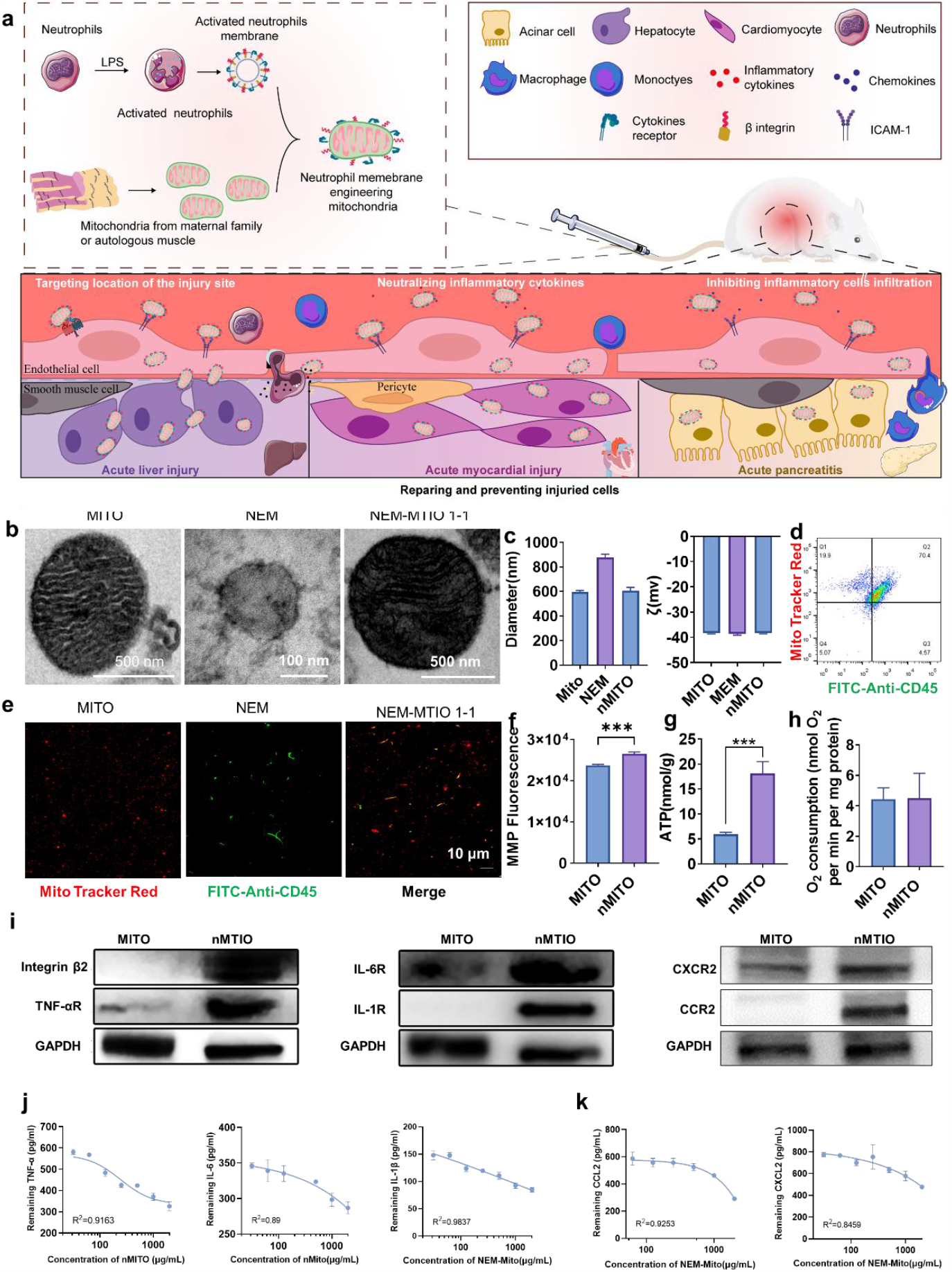
Preparation and Characterization of Neutrophil Membrane Fused Mitochondria (nMITO). **a**. Schematic representation of nMITO designed with organ injury targeting ability, broad-spectrum anti-inflammatory activity, and cell repairing effect. **b**. Representative transmission electron microscopy images of free mitochondria (MITO), neutrophil membrane (NEM), and nMITO. Uranyl acetate staining was applied. Scale bar, 500 nm. **c**. Dynamic light scattering measurements of MITO, NEM, and nMITO hydrodynamic size (diameter) and zeta potential (ζ). **d**. CD45 expression on MitoTracker^®^ Red-labeled nMITO obtained at NEM-MITO ratio of 1/1 according to flow cytometry assay. **e**. CD45 expression on MitoTracker^®^ Red-labeled nMITO obtained at NEM-MITO ratio of 1/1 observed by confocal laser scanning microscopy (CLSM). Scale bar represents 10 μm. **f**. Mitochondrial membrane potential (MMP) indicated by the fluorescence of MitoTracker^®^ Red FM. **g**. Adenosine triphosphate (ATP) synthesis activity of mitochondria. **h**. Oxygen consumption of mitochondria. **i**. Characteristic protein bands of MITO and nMITO resolved by western blotting. Data presented as mean ± s.d. In all datasets, n = 3 independent experiments using the same batch of MITO and nMITO. *, **, and *** denote statistical significance at p<0.05, p<0.01, and p<0.001, respectively. **j-k**. Neutralization curve of nMITO on inflammatory cytokines (j) and chemokines (k). Data presented as mean ± s.d. *, **, and *** denote statistical significance at p<0.05, p<0.01, and p<0.001, respectively.

## Results

### Immuno-engineered mitochondria preparation and characterization

To synthesize nMITO, mitochondria (MITO) and activated neutrophil plasma membrane (NEM) were respectively obtained and purified from mouse heart tissue (Supplementary Fig. 1) and neutrophils (Supplementary Fig. 2, 3). Then, MITO and NEM were successfully fused as nMITO by the fusing of membrane under ultrasound stimulation in ice bath at the NEM: MITO ratio of 1:1, after a screening of the preparation method by the CD45 expression on nMITO (Supplementary Fig. 4-5). Compared with NEM and MITO, nMITO maintained similar morphology, particle size and zeta potential characteristics as MITO (Fig.1b, c). At the same time, nMITO inherited the marker protein CD45 from NEM, and the expression of it increased with the proportion of NEM (Fig. 1d, e and Supplementary Fig. 6). Importantly, this modification did not change the mitochondrial activity of nMITO. On the contrary, it showed higher mitochondrial membrane potential (MMP) and more stable ATP production activity than MITO under the same oxygen consumption level (Fig. 1f-h, Supplementary Fig. 7). Due to the importance of mitochondrial membrane integrity for mitochondrial function, the improved mitochondrial activity may be contributed by strengthening the outer membrane of mitochondria, which need further exploration.

Since cell adhesion receptor from NEM (β-Integrin),chemokine receptors (CXCR2, CCR2) and cytokine receptors (IL-6R, TNF-αR、 IL-1 βR) were also successfully assigned to nMITO (Fig. 1i), we further observed the neutralizing effect of nMITO on inflammatory cytokines, the inhibiting effect on the migration of inflammatory cells and the inhibiting effect on neutrophil adhesion to inflamed endothelial cells. As shown in Fig. 1j, nMITO could significantly neutralize the main inflammatory cytokines in vitro, including TNF-α, IL-1β and IL-6, which are the main cytokines driving the process of inflammation cascade in AOIs^20-23^. In addition, chemokines (such as CXCL-2 and MCP-1) that driving inflammatory cells such as macrophages and neutrophils to migrate to injured tissues^24, 25^ were both neutralized by nMITO (Fig. 1k).

Overall, immuno-engineered mitochondria with retained mitochondrial activity and the neutralizing effect of inflammation cytokines and chemokines were successfully obtained through a rapid preparation method within 30min.

### The targeting ability inherited from neutrophil membrane

In AOIs, neutrophils are known to engage in receptor-mediated adhesion with cytokine-activated vascular endothelial cell and injured tissue cells^26^. Here, nMITO were fluorescently labelled with MitoTracker^®^ Red and added to H_2_O_2_ activated HUVECs (human umbilical vein endothelial cells) or un-activated HUVEC, on which the expression of intercellular activation molecule 1 (ICAM-1) was much higher than un-activated HUVECs (Fig. 2a). Either activated or un-activated HUVECs, when incubated with nMITO, both showed a significant increase in MitoTracker^®^ Red fluorescence compared to MITO (Fig. 2b-c, Supplementary Fig. 8). Moreover, the flow results showed that this difference had nothing to do with the concentration of mitochondria. At each mitochondrial concentration, the average fluorescence intensity of HEVEC incubated with nMITO was higher than that with MITO (Supplementary Fig. 9). Next, we further evaluated the ability of nMITO to distinguish normal vascular endothelium and inflamed vascular endothelium in injuring lesions. Due to the high expression of ICAM-1 on the injured cell surface, we found that the uptake of nMITO by activated HUVEC was much higher than that of naive cells (without H_2_O_2_ activation) (Figure 2c, Supplementary Fig. 8).

**Figure 2.**
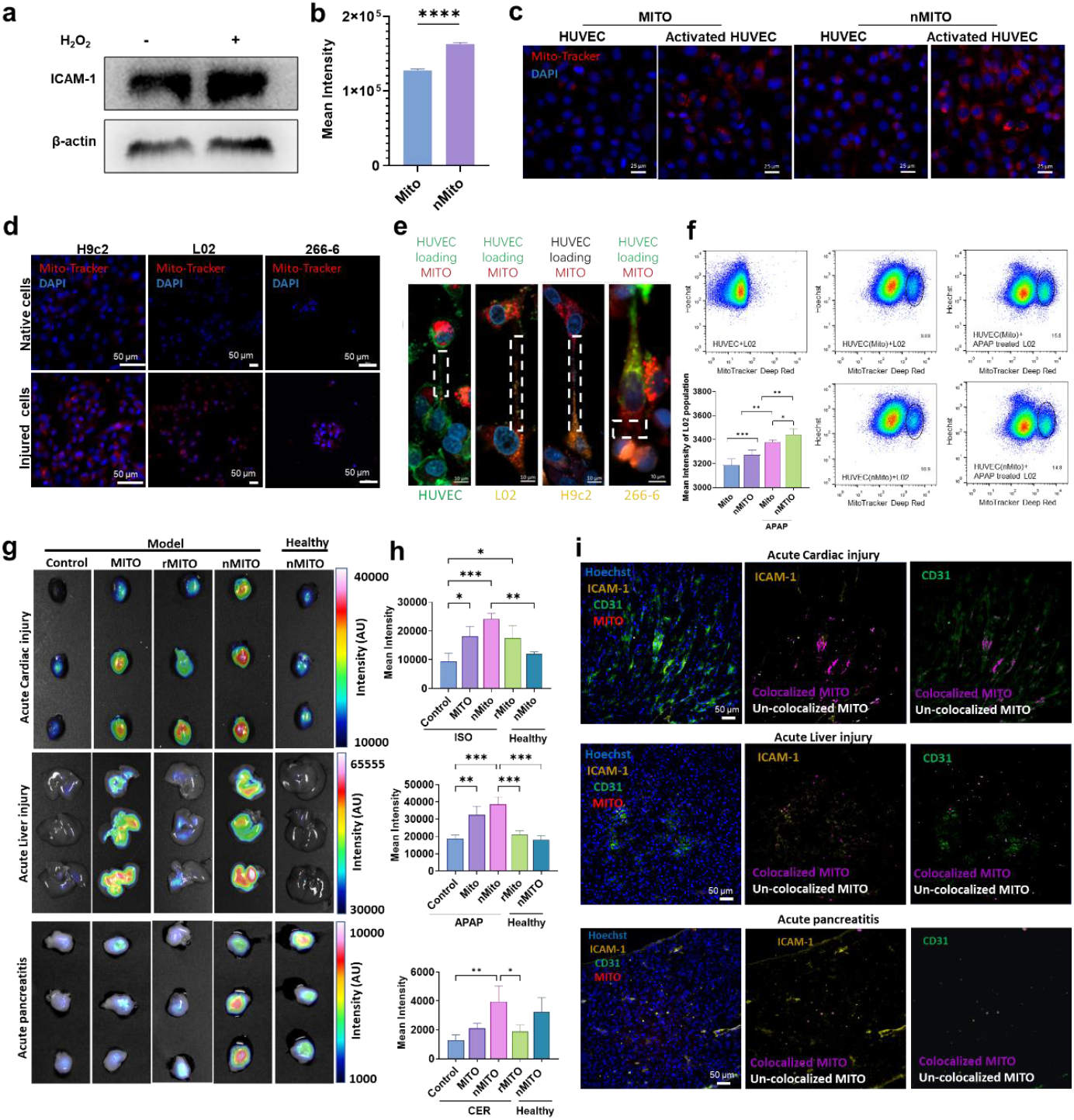
The Targeting Ability and Inflammation Neutralizing Effect of nMITO Inherited from Neutrophil Membrane. **a**. ICAM-1 expression on HUVECs. **b**. Uptake of nMITO and natural mitochondria (MITO) by activated HUVECs examined by flow cytometry assay. **c**. Uptake of nMITO and MITO by activated or naive HUVECs examined by confocal laser scanning microscopy (CLSM). Scale bar represents 25 μm. **d**. Uptake of nMITO and MITO by activated or naive tissue cells examined by CLSM. Scale bar represents 50 μm. **e**. Delivery of ingested mitochondria labeled with MitoTracker^®^ Red from HUVECs to other cells observed by CLSM. Scale bar represents 10 μm. The white dashed box refers to tunneling nanotube. **f**. Delivery of ingested mitochondria from HUVECs to L02 cells monitored by flow cytometry assay. **g-h**. Fluorescent images (g) and quantitative fluorescent intensity (h) of injured heart, liver, and pancreas from mice bearing isoproterenol (ISO)-induced acute cardiac injury, acetaminophen (APAP)-induced acute liver injury, and cerlein (CER)-induced acute pancreatitis. **i**. Location of mitochondria in injured heart, liver, and pancreas from mice bearing ISO-induced acute cardiac injury, APAP-induced acute liver injury, and CER-induced acute pancreatitis. Scale bar represents 50 μm.

Studies have shown that myocardial cells^27^, liver cells^28^ and acinar cells^29^ can also overexpress ICAM-1 on their surfaces in the inflammatory environment or in the case of cell damage, thus being recognized by β Integrin. Therefore, we further observed the difference in uptake of nMITO by damaged cells and healthy cells, and found that the efficiency of nMITO uptake by damaged H9c2 (rat heart ventricular myoblast cell line), L02 (human hepatocyte cell line) and primary acinar cells were all much higher than that of healthy cells (Fig. 2d, Supplementary Fig. 10-12). These results demonstrate the ability of nMITO to target inflamed vascular endothelial and injured tissue cells derived from the fused neutrophil membrane on its surface. The binding is probably attributed to specific interactions between β-integrin on the neutrophil membrane and the ICAM-1 overexpressed on activated HUVECs and injured tissue cells ^30^. As well known, the size of mitochondria is more than 500 nm, which means that it is not easy for mitochondria to directly pass through the endothelial cell leakage into injured tissue^31^. As transfer of mitochondria via tunneling nanotubes (TNTs) is widespread among cells^10, 32^, we found that the mitochondria taken up by activated endothelial cells could be transmitted to the surrounding injured tissue cells, and further transmitted between injured cells via TNTs (Fig. 2e, Supplementary Fig. 13). In addition, the flow results proved that the efficiency of endothelial cells transmitting mitochondria to injured cells is also higher than that to normal cells (Fig. 2f).

To study the ability of nMITO to target and penetrate injured tissues, the mouse models of acute myocardial injury, acute liver injury, and acute pancreatitis were established, and the targeting capability of MitoTracker^®^ Deep Red labelled nMITO, red blood cell membrane coated mitochondria (rMITO) or MITO was assessed. We found that the MitoTracker^®^ Deep Red fluorescence of injured heart, injured liver and injured pancreas in nMITO treated mice were much higher than that in MITO or rMITO treated mice (Fig.4g, h), while there was no significant difference of MitoTracker^®^ Deep Red fluorescence distribution in non-target organs of various AOIs between nMITO, MITO and rMITO (Supplementary Figs. 14–16). In addition, the MitoTracker^®^ Deep Red of injured heart, injured liver and injured pancreas in nMITO treated mice were much higher than that of corresponding tissues in healthy mice treated with nMITO (Fig.4g, h). Further observation by CLSM showed that nMITO were mainly distributed in ICAM-1 high-expressing vascular endothelial cells and tissue cells in section of injured heart, liver and pancreas (Fig.4i, Supplementary Figs. 17–22). The results of organ distribution indicated that nMITO was conducive for optimizing and promoting the distribution of MITO in the injured tissues and injured cells.

Overall, these *in vitro* and *ex vivo* results demonstrate the ability of nMITO to target inflamed endothelial cells and then be further transferred to injured tissue cells.

### The cell repairing activity of nMITO inherited from mitochondria

Numerous studies have demonstrated that exogenous mitochondria can effectively repair damaged mitochondria in injured cells, thereby restoring ATP production, reducing ROS generation, and preventing cell death^33^. To evaluate the cell repair activity of mitochondria, we established cell models of myocardial cell injury, hepatocyte injury, and acinar cell injury and compared the effects of various mitochondrial preparations on these models. In the isoproterenol (ISO)-induced myocardial cell injury model (Fig. 3a, Supplementary Fig. 23), the MMP of damaged cardiomyocytes significantly increased after treatment with MITO and nMITO (Fig.3b, c), and ATP production was significantly restored (Fig. 3d). Notably, nMITO exhibited superior enhancement of MMP and ATP production at most concentrations (Supplementary Fig. 24). Treatment with nMITO and MITO also ameliorated the hypertrophic morphology of myocardial cells (Fig. 3e). Similar results were observed in the acetaminophen (APAP)-induced hepatocyte injury model (Fig. 3f, Supplementary Fig. 25), where treatment with MITO and nMITO significantly increased MMP (Fig.3g, h), restored ATP production (Fig. 3i), and reduced ROS levels (Fig. 3j). Analysis of the markers of hepatocyte injury, including alanine aminotransferase (ALT) and aspartate aminotransferase (AST), indicated that treatment with MITO and nMITO significantly ameliorated hepatocyte injury (Fig. 3k, l), with nMITO showing superior efficacy at most concentrations (Supplementary Fig. 26). We also evaluated the repair effects of mitochondrial preparations on cerulein (CER) and sodium taurocholate (NaT)-induced injury in primary pancreatic acinar cells (Fig. 3m). In the CER and NaT-induced pancreatic primary acinar cells injury model, MITO and nMITO significantly rescued the decreased MMP, with nMITO exhibiting a stronger effect than MITO (Fig.3n-p, Supplementary Fig.27, 28). Treatment with MITO and nMITO also inhibited the opening of mitochondrial permeability transition pore (MPTP), resulting in lower concentrations of Ca^2+^ detected in the treated cells than in the model cells (Fig.3n, o, Supplementary Fig.27, 28). Consequently, the necrosis of acinar cells was significantly inhibited by nMITO (Fig.3n, o, q, Supplementary Fig.27, 28).

**Figure 3.**
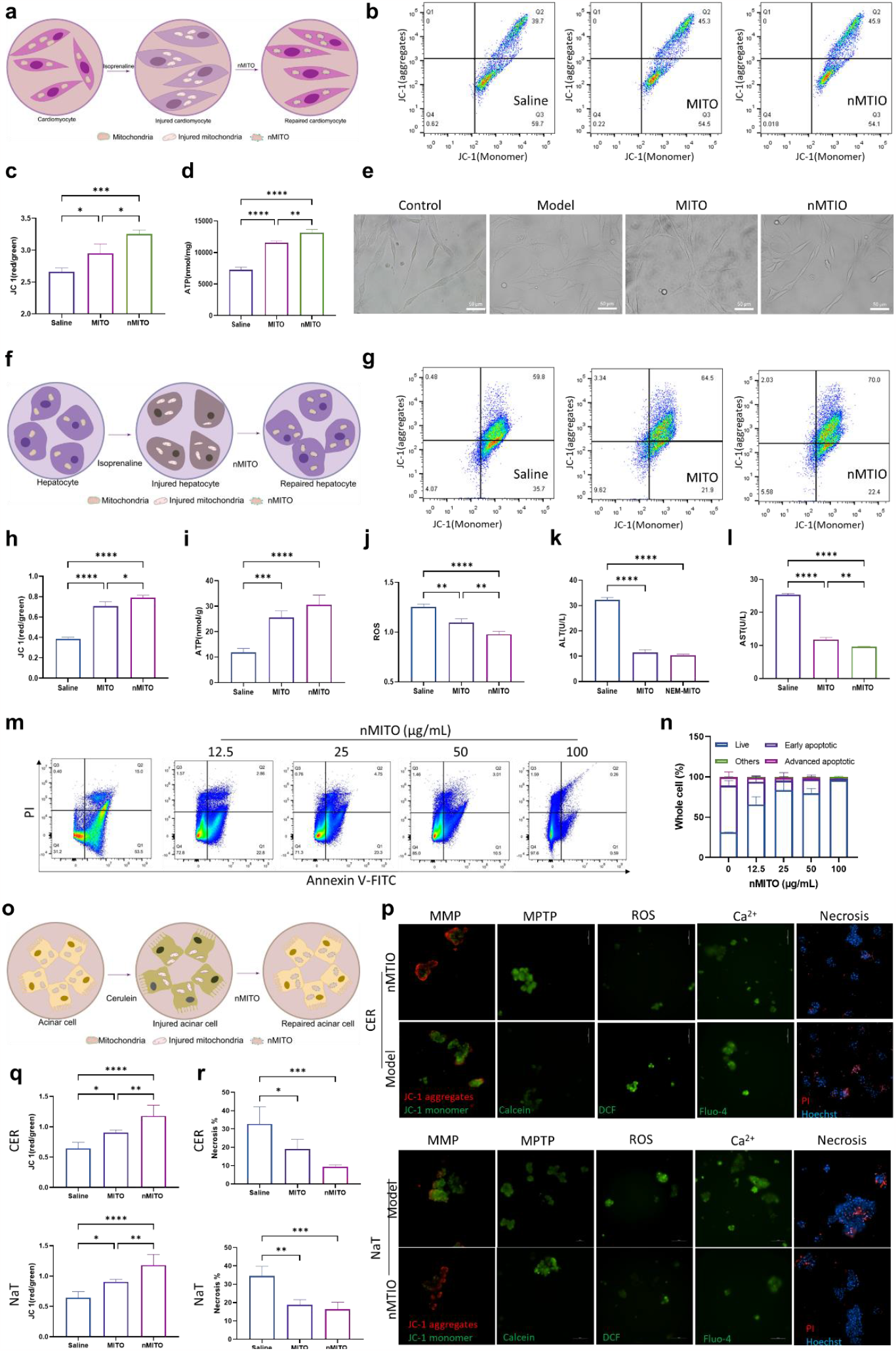
*In Vitro* Cell Repairing Activity of nMITO against Various Injured Cells. **a-e. *In vitro* cell repairing activity against injured H9c2 myocardial cells. a**. Schematic diagram of nMITO repairing H9c2 myocardial cells stimulated by isoproterenol (ISO); **b**. Flow cytometry measurement of JC-fluorescence in H9c2 cells treated with saline, MITO or nMITO; **c**. Calculation of mitochondrial membrane potential of H9c2 cells treated with saline, MITO or nMITO, based on JC-1 Red/Green signal intensity; **d**. ATP synthesis levels of H9c2 cells treated with saline, MITO or nMITO. **e**. Microscopic bright field images of H9c2 cells, indicating myocardial cell hypertrophy. Scale bar = 50 μm. **f-i. *In vitro* cell repairing activity against injured L02 hepatocyte cells. f**. Schematic diagram of nMITO repairing L02 hepatocyte cells injured by APAP; **g**. Flow cytometry measurement of JC-fluorescence in L02 cells treated with saline, MITO or nMITO; **h**. Calculation of mitochondrial membrane potential of L02 cells treated with saline, MITO or nMITO, based on JC-1 Red/Green signal intensity; **i-l**. ATP synthesis (i), reactive oxygen species (ROS) (j), alanine aminotransferase (ALT) (k), and aspartate aminotransferase (AST) (l) levels in APAP-injured L02 cells treated with saline, MITO or nMITO. **m, n**. Flow cytometry and quantitative results of L02 cell apoptosis and necrosis. **o-r. *In vitro* cell repairing activity against injured primary acinar cells. o**. Schematic diagram of nMITO repairing CER or NaT-induced primary acinar cell injury; **p**. The determination of mitochondrial membrane potential (MMP), mitochondrial permeability transition pore (MPTP) opening, ROS, relative Ca^2+^ intensity and necrosis ratio of CER or NaT treated acinar cells after receive saline or nMITO. **q**. The calculation of MMP, based on JC-1 Red/Green signal intensity of CER or NaT treated acinar cells. **r**. The calculation of necrosis ratio of CER or NaT treated acinar cells. Data presented as mean ± s.d. *, **, and *** indicate statistical significance at p<0.05, p<0.01, and p<0.001, respectively.

These results suggest that nMITO not only inherits the cell repair activity from MITO, but further enhance this cell repair activity, which may be related to the fact that injured cells have better uptake of nMITO than MITO.

### The therapy effect of nMITO for acute cardiac injury

nMITO has shown potential in repairing injured cardiomyocytes and regulating inflammation in vitro. Mitochondrial dysfunction and inflammation are known to be involved in the pathogenesis of ISO-induced myocardial injury^34, 35^. To further investigate the pharmacological implications of nMITO in vivo, we examined the putative effect of nMITO on ISO-induced myocardial injury (Fig. 4a). Treatment with nMITO significantly increased ATP and mitochondrial membrane potential (MMP) in the heart tissue of mice, especially at doses of 0.5 mg/kg and 1 mg/kg, which were even higher than those in healthy mice (Fig. 4b, c, Supplementary Fig. 29). nMITO treatment at doses of 0.25 to 1 mg/kg also ameliorated cardiac hypertrophy induced by ISO-induced myocardial injury (Fig. 4d). Echocardiography was used to assess cardiac function and the results showed that the ejection fraction (EF) of mice treated with nMITO at doses of 0.25 or 1 mg/kg was significantly higher than that in the model group, while the fractional shortening (FS) of mice treated with nMITO at doses of 0.25 to 1 mg/kg were all significantly higher than in the model group (Fig. 4d, Supplementary Fig. 30). Furthermore, the heart weight/body weight (HW/BW) of mice treated with nMITO at doses of 0.25 and 1 mg/kg were significantly lower than in the model group treated with saline, while the heart weight/tibial length (HW/TL) and the left ventricular mass index (LVMI) of mice treated with nMITO at doses of 0.25 to 1 mg/kg were all significantly lower than in the model group treated with saline (Fig. 4e-g).

**Figure 4:**
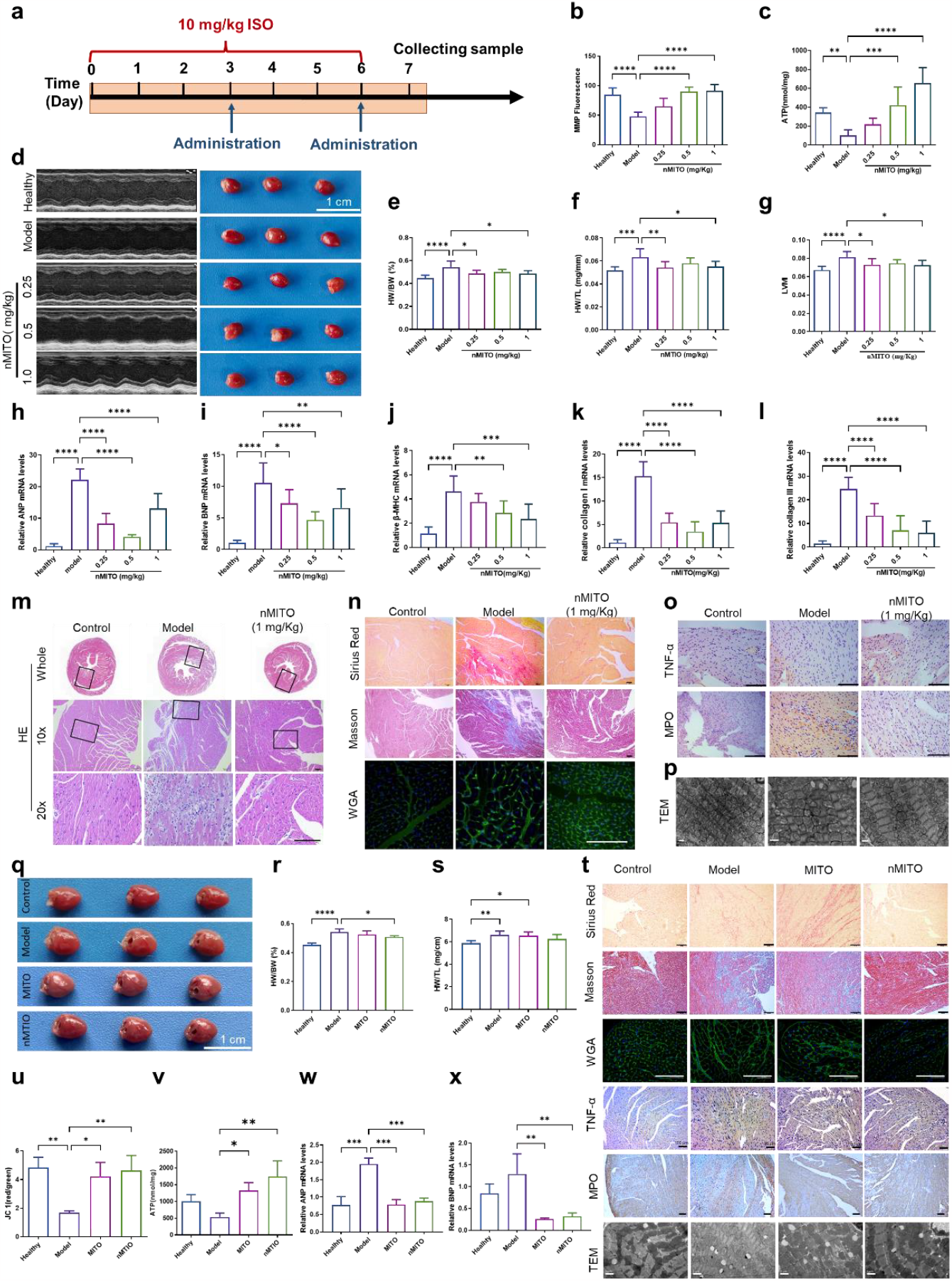
Therapeutic Effect of nMITO in Isoproterenol (ISO)-Mediated Myocardial Injury Animal Model. **a**. Model establishment and administration time. **b-p. n**MITO treatment of myocardial injury effectiveness and dose screening. **b**,**c**. Evaluation of mitochondrial membrane potential (b) and ATP synthesis (c) of cardiomyocytes derived from mice after treatment. **d**. Cardiac ultrasound imaging and general cardiac appearance of mice after treatment. **e-g**. Improvement in the ratio of heart weight to mouse body weight (HW/BW), the ratio of heart weight to tibial length (HW/TL), and the left ventricular myocardial weight (LVM) after treatment. **h**,**i**. Expression of central atrial natriuretic peptide (ANP) and brain natriuretic peptide (BNP) mRNA in heart tissue of mice after treatment. **j-l**. mRNA expression levels of mouse cardiac myosin heavy chain subunit (β-MHC), type I collagen, and type III collagen after different doses of nMITO treatment. **m-o**. Results of hematoxylin-eosin (H&E) staining (m), fibrosis expression (n), and inflammatory infiltration (o) in the myocardium of mice after treatment. **p**. Transmission electron microscope (TEM) observation of mitochondria in myocardial tissue of mice after treatment. **q-v**. Comparison of the therapeutic effect of nMITO versus natural mitochondria (MITO) on myocardial injury. **q**. General appearance of mouse heart after treatment. **s**. Improvement in HW/BW and HW/TL ratios of mice after treatment. **t**. Immunohistochemical results of expression of cardiac fibrosis and inflammatory infiltration in mice after treatment. **u**,**v**. Effect of different preparations on mitochondrial membrane potential and ATP synthesis of mouse cardiomyocytes. **w**,**x**. mRNA expression of ANP and BNP in heart tissue of mice after treatment. Scale bars in all H&E staining, immunohistochemical, and immunofluorescence images are 100 μm. Data presented as mean ± s.d. Scale bars in TEM images are 1 μm. *, **, and *** denote statistical significance at p<0.05, p<0.01, and p<0.001, respectively.

Atrial and ventricular distension and neurohumoral stimuli can stimulate the secretion of natriuretic peptide hormones by the heart, such as atrial natriuretic peptide (ANP) and brain natriuretic peptide (BNP). The mRNA levels of ANP and BNP in ISO-treated mice were both downregulated after nMITO treatment (Fig. 4h, i). Additionally, nMITO at all concentrations significantly reduced the mRNA levels of fibrosis-related proteins such as β-MHC, collagen I, and collagen III (Fig. 4j-l). Hematoxylin & eosin (H&E) staining of heart sections showed that nMITO significantly reduced hypertrophy of myocardial tissue, reduced inflammatory cell infiltration, and reduced the degree of fibrosis in a dose-dependent manner (Fig. 4m). Further immunohistochemistry staining of heart sections showed that nMITO at a dose of 1 mg/kg could ameliorate ISO-induced myocardial fibrosis36, and decrease TNF-α and MPO infiltration in the injured heart (Fig. 4n, o). Compared with the swollen mitochondria in the model, the mitochondria in the cardiomyocytes of mice treated with nMITO were significantly restored, as observed by transmission electron microscopy of heart sections (Fig. 4p). These results suggest that nMITO has therapeutic effects on acute myocardial injury at different doses, with the therapeutic effect being the highest at a dose of 1 mg/kg (Supplementary Fig. 31).

Then, we determined the optimal mitochondrial concentration of nMITO to be 1 mg/Kg and compared its therapeutic effect to that of MITO. In terms of cardiac size, nMITO was superior to MITO in ameliorating myocardial hypertrophy (Fig. 4q). The HW/BW, HW/TL, and LVMI of mice treated with nMITO were all significantly lower than those treated with MITO (Fig. 4r, s). H&E and immunohistochemistry staining of heart sections showed that nMITO reduced myocardial fibrosis, TNF-α, and MPO infiltration more effectively than MITO (Supplementary Fig.32, Fig. 4t). Additionally, nMITO-treated myocardial tissue had more healthy mitochondria compared to MITO-treated myocardial tissue (Fig. 4t). The ATP and MMP levels in the heart tissue of mice treated with nMITO were also higher than those treated with MITO (Fig. 4u, v, Supplementary Fig.33). Furthermore, the mRNA levels of ANP and BNP were lower in nMITO-treated mice compared to MITO-treated mice (Fig. 4w, x). These findings suggest that nMITO, administered at a dose of 1 mg/kg, can significantly attenuate acute myocardial injury induced by ISO. Moreover, compared to MITO, nMITO has distinct advantages in regulating inflammation associated with myocardial injury.

### The therapy effect of nMITO for acute liver injury

Acetaminophen (APAP) is one of the most commonly reported drugs causing drug-induced liver injury and is the leading cause of acute liver failure, accounting for approximately 20% of liver transplantation cases^37^. Herein, APAP induced acute liver injury was employed to evaluate the therapy effect of nMITO for acute liver injury (Supplementary Fig.34). The TEM results showed that the number of mitochondria in injured hepatocytes was significantly increased in mice treated with MITO and nMITO (Fig. 5a). Furthermore, the ATP and ROS levels in mitochondria derived from the liver of nMITO or MITO-treated mice were respectively higher and lower than those in the saline-treated group (Fig. 5b, c). Importantly, nMITO demonstrated more systemic regulation of inflammation than MITO, including the systemic and local inflammation. The serum cytokines, including TNF-α, IL-10, and IL-12, were all significantly decreased to levels similar to those of healthy mice after nMITO treatment (Fig. 5d, f). The immunohistochemical results of liver tissue showed that the cytokines highly expressed in the APAP-injured liver tissue and the infiltration of macrophages and neutrophils were also significantly alleviated in nMITO-treated mice (Fig. 5g, h). Additionally, the concentration of ALT and AST in the serum of mice treated with nMITO was further reduced compared to mice treated with MITO (Fig. 5i, j). H&E staining results also suggested that APAP-mediated liver injury was significantly ameliorated in mice treated with nMITO, while there was still obvious cell necrosis in the liver tissue of mice treated with MITO (Fig. 5k). These findings indicate that nMITO has a stronger therapeutic effect on acute liver injury mediated by APAP than MITO through the recovery of mitochondrial activity and regulation of inflammation.

**Figure 5.**
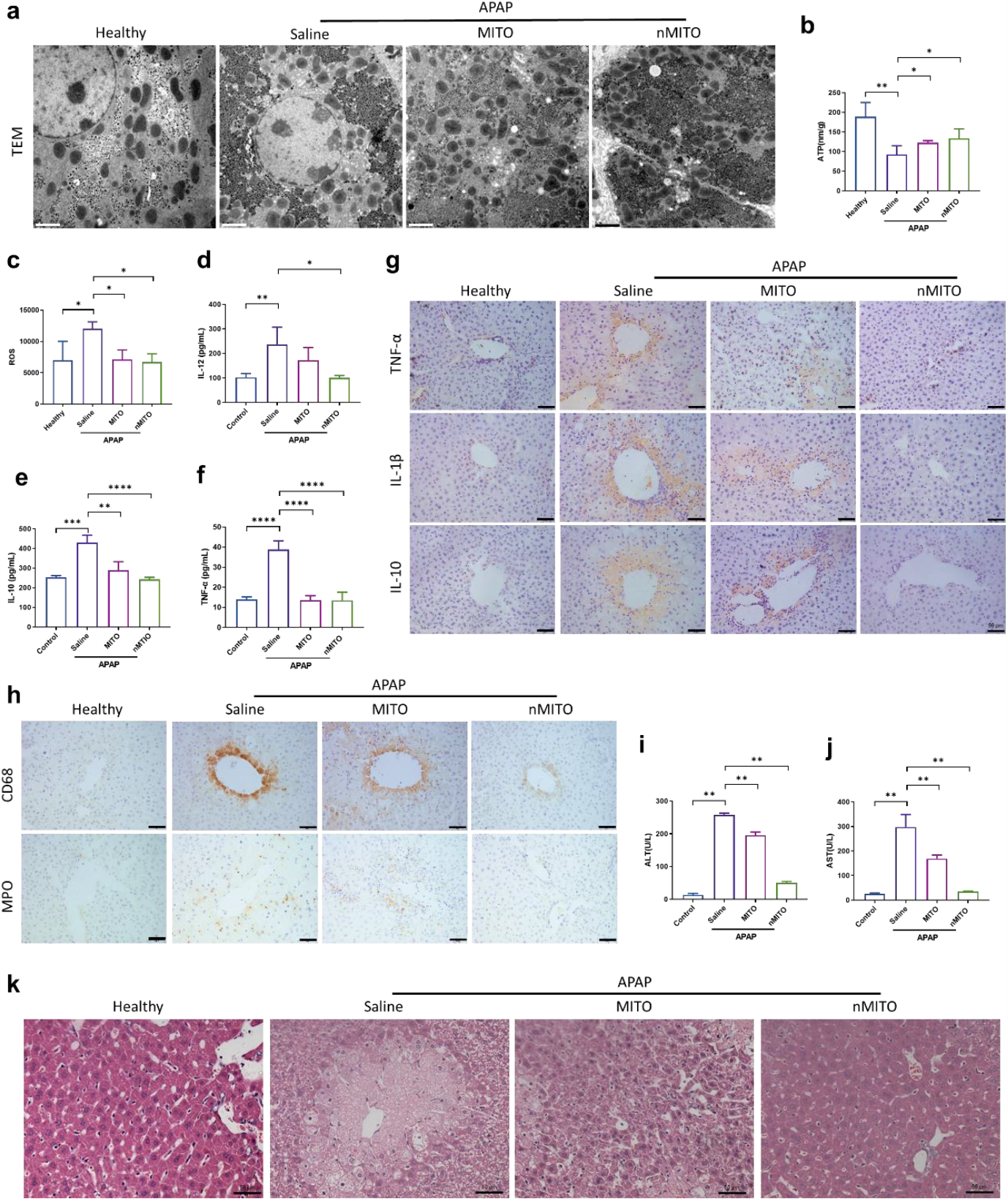
Evaluation of nMITO on the therapeutic effect of acetaminophen (APAP)-mediated acute liver injury. **a**. Transmission electron microscopy (TEM) images of liver tissue mitochondria after treatment with different preparations. **b**,**c**. Evaluation of ATP and ROS activities of hepatocytes after treatment with different preparations. **d-f**. Enzyme-linked immunosorbent assay (ELISA) test results of the infiltration of inflammatory factors in liver tissue after treatment with different preparations. **g**. Immunohistochemical results of inflammatory cytokines infiltration in liver tissue after treatment with different preparations. **h**. Infiltration of macrophages (CD68) and neutrophils (MPO) in liver tissue shown by immunohistochemistry after treatment with different preparations. **i-j**. Results of liver function indexes such as ALT and AST after treatment with different preparations. **k**. H&E staining results of liver tissue after treatment with different preparations. Scale bars in all H&E staining and immunohistochemical images are 50 μm. Data presented as mean ± standard deviation (s.d.). Scale bars in TEM images are 2 μm. *, **, and *** indicate p<0.05, p<0.01, and p<0.001, respectively.

### The therapy effect of nMITO for acute pancreatitis

To investigate the therapeutic potential of nMITO on acute pancreatitis, we tested its effect on cerulein-induced acute pancreatitis (CER-AP) and sodium taurocholate-induced severe acute pancreatitis (NaT-AP)^38^. CER injections caused typical histopathological features of acute pancreatitis, characterized by edema, inflammatory cell infiltration, and pancreatic acinar cell necrosis (Fig. 6a). nMITO significantly enhanced the therapeutic effects of MITO against CER-AP. As shown by histological examination and histopathology scores^39^, nMITO markedly ameliorated CER-induced acinar cell necrosis, pancreatic edema, and infiltration of inflammatory cells, resulting in lower histological scores(Figure 6b,c Supplementary Fig.35). TEM revealed that the pancreatic tissue of mice treated with nMITO has healthier mitochondrial structure than those treated with MITO (Fig. 6b). Additionally, nMITO exhibited a stronger inhibitory effect than MITO on the elevation of serum lipase, amylase, and trypsin induced by pancreatic injury (Fig. 6d-f). Due to the decreased concentration of serum MCP-1 (Fig.6g), nMITO also demonstrated superior inhibition of macrophage in pancreatic tissue (Fig. 6b). Moreover, nMITO significantly reduced the elevated MPO concentration in pancreatic tissue induced by CER (Fig.6h) and the serum concentration of TNF-α in systemic inflammation (Fig. 6i). Notably, these mitochondria had a smaller size than those in healthy pancreatic tissue, likely due to their origin from cardiac tissue (Fig. 6j). These results indicate that nMITO can effectively inhibit the pathological progression of CER-AP through mitochondrial delivery and inflammation regulation.

**Figure 6.**
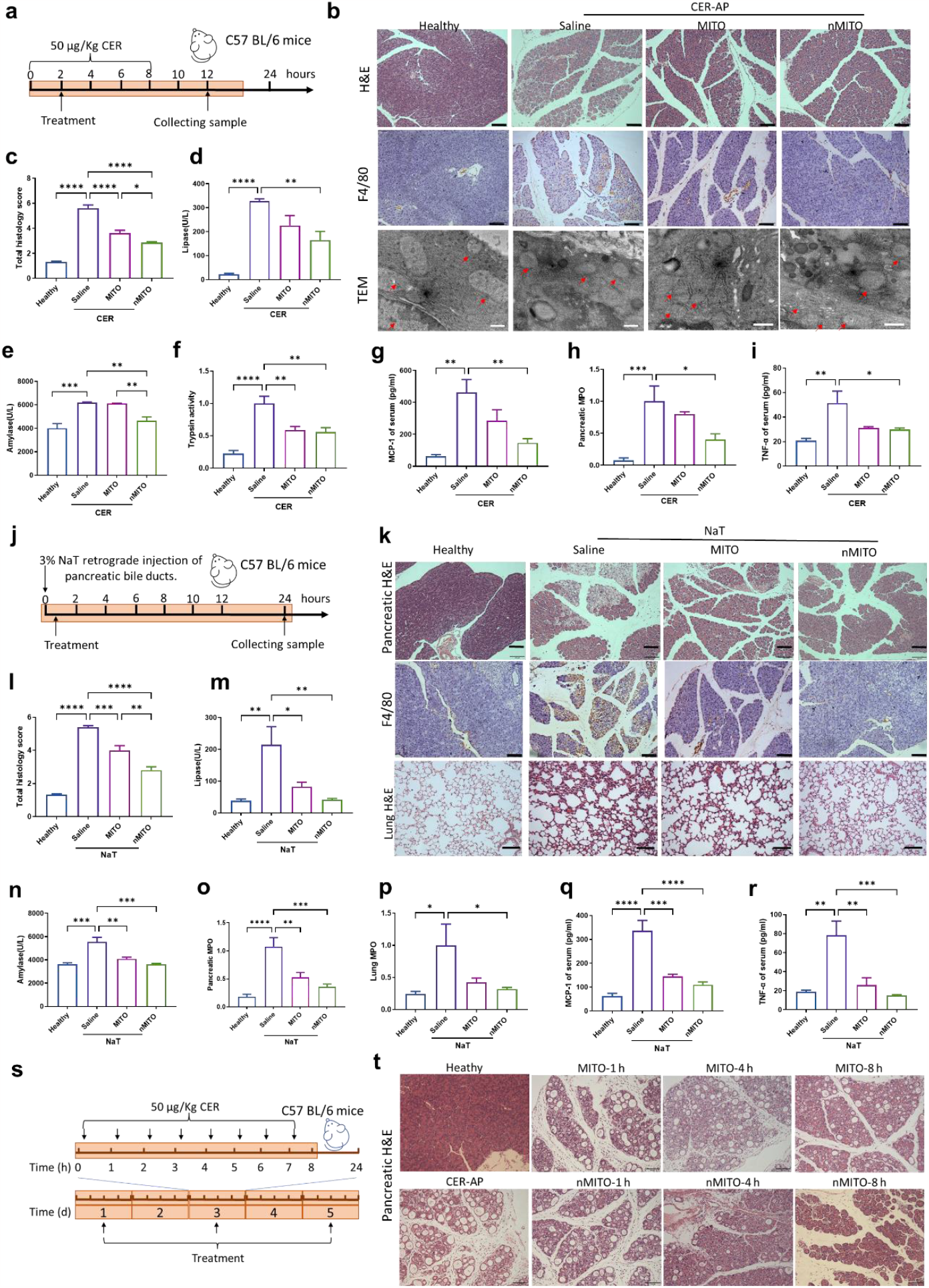
Therapeutic Effect of nMITO in Mouse Models of Acute Pancreatitis. **a-i**. Evaluation of the therapeutic effect of nMITO on cerulein-mediated acute pancreatitis model (CER-AP). **a**. Model establishment and administration time; **b-c**. H&E sections, F4/80 staining, TEM images (b), and total pathological scores (c) of mouse pancreas after treatment with different mitochondrial preparations; **d-f**. Concentration of serum lipase (d), amylase (e) and trypsin (f) in mice after treatment with different mitochondrial preparations; **g**. MCP-1 concentration in mouse serum; **g**. MPO activities in mouse pancreas; **i**. TNF-α concentration in mouse serum. **j-k**. Evaluation of the therapeutic effect of nMITO on the model of severe acute pancreatitis mediated by sodium taurocholate (NaT-AP); **j**. Model establishment and administration time; **k**. H&E sections and F4/80 staining images of mouse pancreas, and H&E section image of mouse lung. **l**. pathological scoresof mouse pancreas after treatment with different mitochondrial preparations; **m-n**. Concentration of serum lipase (m) and amylase (n) in mice after treatment with different mitochondrial preparations; **o**. MPO activities in mouse pancreas; **p**. MCP-1 concentration in mouse serum; **q**. TNF-α concentration in mouse serum; **r**. MPO activities in lung. **s, t**. Evaluation of the therapeutic effect of nMITO on the model of severe acute pancreatitis mediated by cerulein (CER-SAP). **s**. Model establishment and administration time; **t**. H&E sections of mouse pancreas after treatment with different mitochondrial preparations. Data presented as mean ± s.d. Scale bars in all H&E staining, immunohistochemical, and immunofluorescence images are 100 μm. Scale bars in TEM images are 0.5 μm. *, **, and *** denote statistical significance at p<0.05, p<0.01, and p<0.001, respectively.

Furthermore, we constructed a severe acute pancreatitis model by intraductal injection of sodium taurocholate (NaT) (Fig.6j). Both MITO and nMITO markedly ameliorated pancreatic histological injury manifesting as reduced pancreatic edema, acinar cell necrosis, neutrophil infiltration as well as macrophage infiltration (Fig.6j). The protective effects of nMITO on pancreatic histopathology including edema, inflammation, necrosis was more significant than that of MITO (Fig. 6l, Supplementary Fig.36). The dramatic elevations of serum lipase (Fig.6m) and amylase (Fig.6n) by NaT were both inhibited at a similar level in nMITO treated mice. Compared to CER-AP, NaT-AP was characterized by a higher degree of macrophage infiltration in pancreatic tissue (Fig.6k). nMITO significantly reduced macrophage infiltration in pancreatic tissue, and further decreased the pancreatic MPO activity compared to MITO, (Fig.6o). Systemic inflammation and acute respiratory distress syndrome are the commonest complications during SAP, the leading cause of lethality in SAP^40^. nMITO significantly reduced the high serum concentration of MCP-1 and TNF-α induced by NaT-AP (Fig. 6 p,q), and significantly decreased the thickness of alveolar walls, lung MPO activity (Fig. 6k,r). In addition, a severe acute pancreatitis model by injecting CER for 5 days, and mitochondrial therapy was performed at 1h, 4h, or 8h after injection of CER on days 1, 3, and 5 (Fig.6s). In the model, a large amount of necrosis, inflammatory infiltration, and duct metaplasia were observed in mouse pancreatic tissue (Fig.6t). However, at 1, 4, and 8 hours after injection of CER, the pancreatic tissue pathology of mice treated with nMITO improved to some extent, with the best therapeutic effect achieved after 8 hours of CER injection (Fig.6t), and the mice received nMITO also showed a more active state than control mice and mice received MITO (Supplementary movie.1)

These data above suggested that nMITO is more potential in ameliorating acute pancreatitis, particularly in ameliorating inflammation during SAP.

## Discussion

We have developed a novel therapeutic strategy for acute organ injuries (AOIs) using immuno-engineered mitochondria (nMITO), which are created by fusing neutrophil membranes with mitochondria. Our findings demonstrate that nMITO have a broad-spectrum blockade of the inflammatory cascade and provide complete mitochondrial function supplementation, enabling simultaneous modulation of the two common mechanisms involved in AOIs. Furthermore, the presence of β-integrin on nMITO enables them to selectively accumulate in injured tissues by binding to its ligand ICAM-1, which is highly expressed in injured endothelial and tissue cells. This feature further amplifies the regulatory effects of nMITO on local inflammation and the repair of injured cells. Importantly, nMITO can address multiple organ injuries that are frequently associated with AOIs, making them a promising non-specific disease-targeted treatment strategy.

For future clinical translation, neutrophils can be collected from patients’ own blood, and the mitochondria can be obtained from their muscle tissue, platelets, or maternal relatives, due to the maternal genetic characteristics of mitochondria^41^. However, scaling up and manufacturing may be challenging due to the use of natural cells and organelles. Additionally, the rapid inactivation of mitochondria is a limitation for clinical application, but recent progress in the extraction, preservation, quality control, and *in vitro* amplification of mitochondria offer a promising outlook for the clinical transformation of nMITO^42-47^. In summary, our immuno-engineered mitochondria represent a potentially valuable therapeutic strategy for a wide range of acute organ injuries, with the prospect of improving clinical outcomes for affected patients.

## Methods

### Animals

All animals used in this study were treated in strict accordance with the protocol approved by the Chongqing University of Technology Ethics Committee (approval number: 202110). SPF male C57BL/6 mice were procured from Hunan SJA Laboratory Animal Co. Ltd. (Chongqing, China). The mice were housed in a controlled environment with a temperature of 23 ± 2°C, suitable humidity, and a 12-hour light-dark cycle. The mice had unlimited access to food and water.

### Isolation of mitochondria

Mitochondria were isolated from C57BL/6J mouse hearts using the Beyotime Tissue Mitochondrial Isolation Kit. The hearts were cut into small pieces, digested using pre-chilled trypsin, and centrifuged at 4 °C and 600 g for 20 seconds. The precipitate was added to the mitochondrial lysate and homogenized on ice. The homogenate was centrifuged at 1,000 g and 4 °C for 5 minutes, and the supernatant was taken and centrifuged at 3,500 g and 4 °C for 10 minutes. The resulting mitochondria were resuspended in a suitable amount of PBS, and the mitochondrial concentration was determined using BCA assay. The mitochondrial ATP and MMP levels were measured using the ATP and MMP assay kits. The size and zeta potential of the mitochondria were measured using dynamic light scattering (DLS, Brookhaven) and observed by transmission electron microscopy (TEM, JEM-1400 Plus Electron Microscope).

### Neutrophil cell membrane extraction

Mouse neutrophils were collected from C57BL/6J mice using a density gradient centrifugation method. After LPS stimulation (100 ng/ml) for 2 hours, the cells were resuspended in ice-cold isolation buffer (containing 225 mM mannitol, 75 mM sucrose, 30 mM Tris-HCl, 0.5 mM EDTA, and 1% (v/v) protease inhibitor) and subjected to sonication at 100 W for 5 minutes (2 seconds sonication, 3 seconds intervals) on ice. The homogenate was then centrifuged at 10,000×g and 4 °C for 10 minutes, the supernatant was collected, freeze-dried, and stored at -80 °C for further use.

### Preparation and characterization of nMITO

Neutrophil cell membranes (NEM) were resuspended in ddH_2_O water and mixed with freshly extracted mitochondria (MITO) at protein ratios of 1:1, 2:1, and 4:1. The mixture was sonicated for 2 minutes at 4 °C. Subsequently, the uncoated NEM was removed by centrifugation at 3,500 g for 10 minutes at 4 °C, and the precipitate was engineered mitochondria (nMITO). The mitochondria were labeled using MitoTracker ^@^ Red CMX Ros, and the activated neutrophil membranes were labeled with FITC-CD45 Monoclonal Antibody. The colocalization of neutrophil membranes and mitochondria was detected using confocal laser scanning microscopy (CLSM, Nikon) and flow cytometry (BD FACS Celesta). The ATP content of MITO and nMITO was measured separately after extraction, and the mitochondrial morphology after NEM coating was observed using TEM. The following antibodies were used to confirm the presence of membrane-associated proteins on nMITO: anti-Integrin β2, anti-TNFR, anti-IL 6R, anti-IL 1R, CXCR2, CCR2 and GAPDH.

### Quantification of cytokine Binding

TNF-α, IL-1β, IL-6, and CXCL2 were mixed with nMITO at different concentrations ranging from 0 to 2 mg/mL. The mixtures were incubated at 37 °C for 2 h and then centrifuged to remove the mitochondria. The cytokine concentration in the supernatant was quantified using ELISA kits. Nonlinear curve fitting was performed using GraphPad Prism 8.

### Cell culture and treatment

H9c2 (rat heart ventricular myoblast cell line), L02 (human normal liver cell line), and 266-6 (mouse acinar pancreatic cell line) cells were cultured in DMEM and RPMI 1640 supplemented with 10% FBS, 100 U/mL penicillin, and 100 μg/mL streptomycin, and incubated at 37 °C in a cell culture incubator with 5% CO_2_.

### Pancreatic acinar cell preparation

Primary pancreatic acinar cells were isolated from the pancreas of male C57BL/6 mice according to previous studies ^48^. Pancreatic tissue was removed immediately after the mice were sacrificed and injected with collagen IV (200 U/mL) in the pancreas, where they were co-incubated for 19 minutes at 37°C. After incubation, cells were isolated from the pancreas by mechanical blowing and filtered through a 100 μm cell filter. The isolated cells were centrifuged at 700 rpm for 2 min to obtain cell pellets, which were resuspended in a DMEM solution containing 1 % bovine serum albumin (BSA), 10% serum, and 2 % antibiotics. The cells were then treated in cell incubators for 12 hours.

### Injured cell preparation

#### HUVEC

The cells were cultured overnight, and then different concentrations of H_2_O_2_ were added to the media for 2 hours to establish the cell damage model. Cell activity was evaluated using CCK-8, and ICAM-1 expression in cells was analyzed using western blot.

#### H9c2

The cells were cultured overnight, and then different concentrations of isoproterenol (ISO) were added to the media for 12, 24, and 48 hours to establish the cell damage model. Cell activity was evaluated using CCK-8, and cellular ATP content and MMP levels were measured to assess cellular mitochondrial function.

#### L02

The cells were cultured overnight, and then different concentrations of APAP were added to the media for 12, 24, and 48 hours to establish the cell damage model. Cell activity was evaluated using CCK-8, and the levels of cellular ATP content were evaluated using commercial kits. The levels of ALT and AST in the supernatant of L02 cells were measured after centrifugation.

#### Primary acinar cells and 266-6 cells

Acinar cells were isolated and incubated with 200 nM CER or 1.5 mM NaT, respectively, for the next 12 hours to establish the cell damage model. Total cell counts were determined using nuclear staining with Hoechst 33342 (50 μg/mL), and the number of necrotic cells was assessed using PI (1 μmol/mL). The necrosis rate was calculated by dividing the number of necrotic cells by the total number of cells. The 266-6 cell damage model was established in the same way as for primary acinar cells.

### *In vitro* repairing of mitochondria on injured cells

#### H9c2

To investigate the potential of mitochondrial repair in injured cells, H9c2 cells were treated with different concentrations of MITO (6.25, 12.5, 25, 50 μg/ml) or nMITO (6.25, 12.5, 25, 50 μg/ml) for 24 h. MMP and ATP content measured by commercial kits.

#### L02

To investigate the potential of mitochondrial repair in injured cells, H9c2 cells were treated with different concentrations of MITO (6.25, 12.5, 25, 50 μg/ml) or nMITO (6.25, 12.5, 25, 50 μg/ml) for 24 h. Cells were collected, and MMP, ATP content, AST, ALT and ROS levels were measured separately using commercial kits.

#### Primary acinar cells

To investigate the potential of mitochondrial repair in injured cells, fresh acinar cells were isolated and treated with 200 nM CER or 1.5 mM NaT respectively. After 30 minutes, different concentrations of MITO or nMTO (6.25, 12.5, 25 μg/ml) were added to each well for the next 12 hours. Nuclear staining with Hoechst 33342 (50 μg/mL) was used for total cell counts, while PI (1 μmol/mL) was used to assess necrotic cell numbers. The MMP, ROS, relative calcium (Ca^2+^) intensity, and MPTP opening were determined using the MMP assay kit, DCFH-DA probes, Fluo-4 AM probe, and MPTP detection kit, respectively. Fluorescent images were captured by a fluorescent microscope and analyzed using Image J software. Additionally, ATP levels were measured using an ATP detection kit.

### Cell uptake

H9c2, L02, and 266-6 cells were cultured overnight in a glass-bottom cell culture dish and stimulated with ISO, APAP, or CER as described earlier. Mito and nMITO, labeled with MitoTracker^®^ Red CMX Ros, were added to the freshly prepared media. After 24 hours of incubation, the cells were washed three times, fixed with paraformaldehyde, and their nuclei were stained with DAPI. The cells were observed and photographed using CLSM (Nikon) or High-content imaging systems (GE IN Cell Analyzer 2500HS). The efficiency of mitochondrial transformation into cells was measured by analyzing the mean fluorescence intensity using flow cytometry.

### The transfer of mitochondria between cells

Mitochondrial transfer between endothelial cells and H9c2, L02, or 266-6 cells was investigated. A model of HUVEC that had been injured was built, and cells were collected after co-incubating with nMITO labeled by MitoTracker^®^ Deep Red for 12 hours. For an additional 12 hours, HUVEC were co-incubated with H9c2, L02, or 266-6 cells, respectively. The transmission of mitochondria between cells was investigated using flow cytometry (ACEA NovoCyte D2060R) and CLSM (Zeiss LSM900).

### Treatment for mice bearing APAP-induced liver injury

#### APAP-induced liver injury

C57BL/6J mice were fasted overnight prior to the administration of 400 mg/kg APAP dissolved in PBS. The mice were then randomly assigned to one of four groups (n = 10 for each group). The first group served as the normal control, while the remaining three groups received i.p. injections of APAP. Two hours after APAP administration, mice in the mitochondria-treated group were injected with a saline solution of either MITO or nMITO (1 mg/kg) via the i.v. route, while mice in the negative control group were injected with an equal volume of saline.

#### Measurements

After 24 hours, all mice were euthanized and serum and liver samples were collected. Commercial kits were used to measure serum ALT and AST activity, ROS levels, and cytokine levels, according to the manufacturer’s protocol. In addition, mouse liver ATP and MMP levels were measured. To observe intracellular mitochondria, mouse liver samples were fixed in 2.5% glutaraldehyde, embedded, and sectioned, and images were captured using TEM (Hitachi H-7650). Tissue sections of the large lobes were also fixed in 4% paraformaldehyde for further analysis.

### Treatment for mice bearing ISO-induced cardiac injury

C57BL/6J mice were fasted overnight before the first administration of ISO. The mice were randomly assigned to five groups (n = 8-10 for each group). The first group served as a normal control. The other four groups received 10 mg/kg ISO (2 mg/mL, 5 mL/kg, dissolved in 0.1% VC solution) subcutaneous injection once daily for 7 days. The nMITO treatment group received 0.25 mg/kg, 0.5 mg/kg, and 1 mg/kg of nMITO (10 mL/kg) intravenous injection (i.v.) via tail veins 12 hours after ISO injection on days 3 and 6. At 24 hours after the ISO injection on day 7, all mice were euthanized, and serum and heart samples were collected.

Mouse cardiac function was evaluated using small animal ultrasonography 12 hours after the second mitochondrial injection treatment. After the heart ultrasonography tests, all mice were weighed, and the body weight, heart weight, and tibia length were measured. The ratios of heart weight to body weight (HW/BW) and heart weight to tibia length (HW/TL) were calculated. Mouse heart ATP and MMP levels were measured using commercial kits, similar to the liver-injured mice. Mouse hearts were fixed immediately with 4% paraformaldehyde and 2.5% glutaraldehyde, respectively, and then embedded for tissue sectioning or TEM. ANP, BNP, β-MHC, collagen I, and collagen III mRNA levels were measured simultaneously.

### Treatment for mice bearing experimental AP

#### NaT-induced AP

Male C57BL/6J mice were fasted overnight before NaT administration. The mice were randomly assigned to four groups (n = 8 for each group). The first group served as the normal control, while the other three groups were induced with AP (NaT-AP) by 3% NaT retrograde injection of pancreatic bile ducts. Prior to the operation, mice were anesthetized with 1.2% tribromoethanol (20 μL/g) and kept warm after the operation. If the mice experienced excessive distress throughout the experiment, they were humanely euthanized. A saline solution of either MITO or nMITO (0.5 mg/kg) was injected intravenously 30 min after operation, while the control group was injected with the same dose of normal saline. Twenty-four hours after the operation, all mice were euthanized, and samples of serum, lung, and pancreatic tissue were collected.

#### CER-induced AP

Acute pancreatitis was induced by 9 hourly intraperitoneal injections of 50 μg/kg CER (CER-AP). The control group was injected with the same dose of normal saline according to the same schedule. The mice were randomly assigned to four groups (n=6). A saline solution of either MITO or nMITO (0.5 mg/kg) was administered at the second dose of CER, while the control group was injected with the same dose of normal saline. Mice were humanely euthanized 12 hours after the first injection of CER, and serum and tissue were collected for further study.

#### Measurements

Serum lipase, amylase, and cytokine levels were measured using commercial kits, following the manufacturer’s protocols. To evaluate pathological changes, pancreatic tissue sections were stained with H&E and examined under a light microscope^49^. Two pathologists made a blinded assessment of edema, inflammatory factor infiltration, and necrosis in the pancreatic sections. The scores ranged from 0 to 3, and the total pathological score was obtained by summing the scores of inflammations, edema, and necrosis (scores range from 0 to 9) ^50^. The infiltration of macrophages in pancreatic tissue was observed by staining with Anti-F4/80 antibody. To observe intracellular mitochondria, mouse pancreas samples were fixed in 2.5% glutaraldehyde, embedded, and sectioned, and images were captured using TEM (Hitachi H-7650). Myeloperoxidase (MPO) activity was determined using the method described in a previous study^51^. To determine the activation of pancreatic trypsin, pancreatic tissues were homogenized in homogenate buffer and centrifuged at 1500 × g for 5 min. The supernatant was collected and reacted with 10 mM Boc-Glu-Ala-Arg-MCA substrate. Finally, the fluorescence was measured and normalized to protein concentration at 100%.

### *In vivo* distribution of nMITO

Fluorescence *in vivo* imaging was performed using an in vivo imaging system (VILBER FUSION FX). C57BL/6J mice were randomly divided into five groups, with 3-4 mice in each group. To induce liver injury and cardiac injury models, MitoTracker^®^ Deep Red labeled MITO, nMITO, and rMITO at 1 mg/kg were injected. Healthy mice were injected with the same amount of nMITO as a control. Two hours after the injections, all mice were subjected to ultrasonography, and their hearts, livers, kidneys, spleens, and lungs were imaged. Heart, liver and pancreas tissue samples from mice with cardiac injury, liver injury and acute pancreatitis respectively, were collected and frozen. The frozen sections (6 μm) of heart, liver and pancreas were cut using a cryomicrotome (LEICA CM1950). The sections were then fixed and stained with FITC anti-CD31 for blood vessels, anti-ICAM1 for damaged cells, and Hoechst 33342 for nuclei. The fluorescence of the sections was observed under a high-content imaging system (GE, IN Cell Analyzer 2500HS).

### Statistical analysis

We used GraphPad Prism version 9.0 for statistical analysis of the data. Comparisons between two groups were accomplished using independent sample t-test, while groups are more than two groups, ordinary one-way analysis of variance (ANOVA) was employed. Results were presented as mean ± SD from at least three independent experiments. A value of P < 0.05 was considered statistically significant.

## Acknowledgments

This study was financially supported by Chongqing Municipal Technology Innovation and Application Development Special Key Project (CSTB2021TIAD-KPX0051), National Natural Science Foundation (82100684), Chongqing medical scientific research project (Joint project of Chongqing Health Commission and Science and Technology Bureau, No. 2022MSXM154), and Postdoctoral Research Foundation of China (2021M700625).

## Author contributions

Xing Zhou, was primarily responsible for the research idea, literature review, research design, and foundation application. Qing Zhang, Hanyi Zhang, and Xuemei Li performed the preparation of engineered mitochondria. Hanyi Zhang, Shengqian Yang and Jinwei Feng conducted *in vitro* and *in vivo* experiments related to myocardial injury. Qing Zhang, Xuemei Li, and Chenglu Hu conducted *in vivo* and *in vitro* experiments related to acute liver injury. Yan Shen, Cheng Dai, Xiuyan Yu, and Zhihua Lin conducted *in vivo* and *in vitro* experiments related to acute pancreatitis. Xing Zhou, Jie Lou and Chengyuan Zhang conducted statistical analysis of data. Zhihua Lin, Xiaohui Li and Xing Zhou supervised the work. Xing Zhou, Qing Zhang, and Yan Shen wrote the manuscript, with contributions from all other authors.

## Supplementary information

Supplementary methods and Supplementary Figs. 1–35.

