## Supplementary Information for "Immuno-Engineered Mitochondria for Efficient Therapy of Acute Organ Injuries via Modulation of Inflammation and Cell Repair"

**Supplementary Methods**

**Materials**

Isoprenaline (MCE, American, HY-B0468); Acetaminophen (RHAWN, China, R018054); CER (Cellmano, China, C0301); Sodium taurocholate (NaT, 86339), LPS (L4391), collagen IV (C9407) purchased from sigma (American); MitoTracker® Deep Red FM (Yeasen, China, 40743ES50); Mito Tracker® Red CMX Ros (M7512), FITC-CD45 Monoclonal Antibody (11-0451-82), IL1RA Polyclonal Antibody (PA5-21776), TNFR2 Recombinant Rabbit Monoclonal Antibody (MA5-32618), IL-6R Polyclonal Antibody (PA5-102425) purchased from ThermoFisher Scientific (American); GAPDH Monoclonal Antibody (0004-1-Ig), FITC Anti-Mouse CD31(FITC-65058) purchased from Proteintech Group (American); Anti-CCR2 Recombinant Rabbit Monoclonal Antibody (ET 1611-65) and Anti-F4/80 antibody (RT1212) purchased from HUABIO (China); Anti-ICAM1 antibody (Abcam, UK, ab222736, ab109361); Rabbit Anti-CXCR2 antibody (Bioss, China, bs-4836R); Integrin β2 Polyclonal Antibody (Immunoway, China, YT5461); MPO Antibody (R&D Systems, American, AF3667); Anti-CD68 Rabbit pAb (GB11067), Propidium iodide (PI, G1021), Phosphate Buffered Saline (G0002-15) purchased from Servicebio (China); Cytokine assay kit purchased from NeoBioscience (China, MC102a.96, EMC001b.96, EMC004.96, EMC122.96, EMC113.96); Mouse Bone Marrow Neutrophil Isolation Kit (Solarbio, China, P8550); Tissue Mitochondrial Isolation Kit (C3606), BCA Protein Assay Kit (P0011), Mitochondrial membrane potential assay kit with JC-1 (C2006), ATP Assay Kit (S0027), Protease inhibitor (P1005), Cell Counting Kit-8 (C0038), Reactive Oxygen Species Assay Kit (S0033S), Hoechst 33258 (C1011), Antifade Mounting Medium with DAPI (P0131), Fluo-4 AM (S1060), MPTP detection kit (C2009S), hematoxylin and eosin (H&E , C0105S) all purchased from Beyotime Biotechnology(China); Alanine aminotransferase (C009-2-1), Aspartate aminotransferase (C010-2-1), Serum lipase (A054-2-1), amylase (C016-1-1) all purchased from Nanjing Jiancheng Institute of Bioengineering (China).

**Isolation of Mitochondria**

Mitochondria were isolated from C57BL/6J mouse hearts using the Beyotime Tissue Mitochondrial Isolation Kit. The hearts were cut into small pieces, digested using pre-chilled trypsin, and centrifuged at 4 °C and 600 g for 20 seconds. The precipitate was added to the mitochondrial lysate and homogenized on ice. The homogenate was centrifuged at 1,000 g and 4 °C for 5 minutes, and the supernatant was taken and centrifuged at 3,500 g and 4 °C for 10 minutes. The resulting mitochondria were resuspended in a suitable amount of PBS, and the mitochondrial concentration was determined using BCA assay. The mitochondrial ATP and membrane potential (MMP) levels were measured using the ATP and JC-1 assay kits. The size and zeta potential of the mitochondria were measured using dynamic light scattering (Brookhaven) and observed by transmission electron microscopy (JEM-1400 Plus Electron Microscope).

**Neutrophil Cell Membrane Extraction**

Mouse neutrophils were collected from the bone marrow of C57BL/6J mice using a density gradient centrifugation method. After LPS stimulation (100 ng/ml) for 2 hours, the cells were resuspended in ice-cold isolation buffer (containing 225 mM mannitol, 75 mM sucrose, 30 mM Tris-HCl, 0.5 mM EDTA, and 1% (v/v) protease inhibitor) and subjected to sonication at 100 W for 5 minutes (2 seconds sonication, 3 seconds intervals) on ice. The homogenate was then centrifuged at 1,000×g and 4 °C for 10 minutes, the supernatant was collected, freeze-dried, and stored at -80 °C for further use.

**Supplementary Figures**

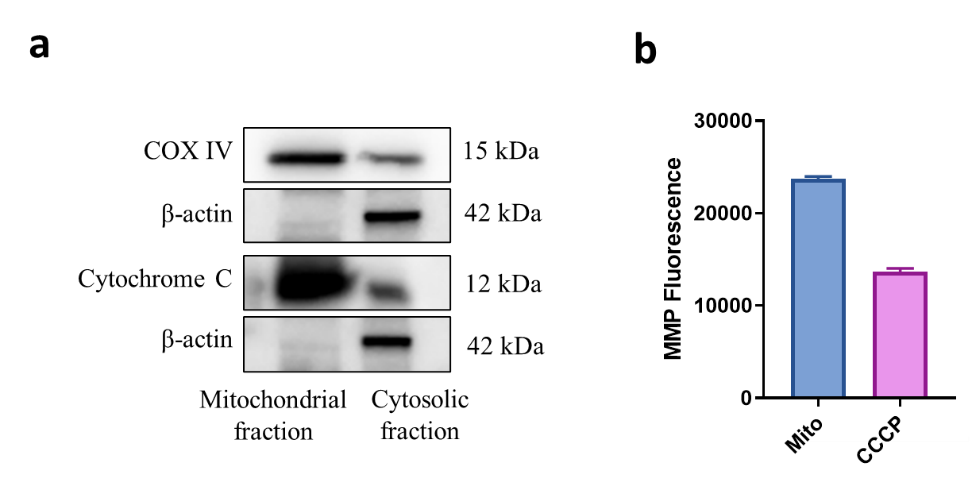

**Supplemental. Figure. 1.** The characterization of isolated mitochondria. a. The representative proteins of mitochondria (COX IV and Cytochrome C), Proliferating Cell Nuclear Antigen (PCNA) and cytoskeleton (β-actin) in mitochondrial fraction and cytosolic fraction. b. The mitochondrial membrane potential (MMP) indicated by the fluorescence of MitoTracker® Red FM.

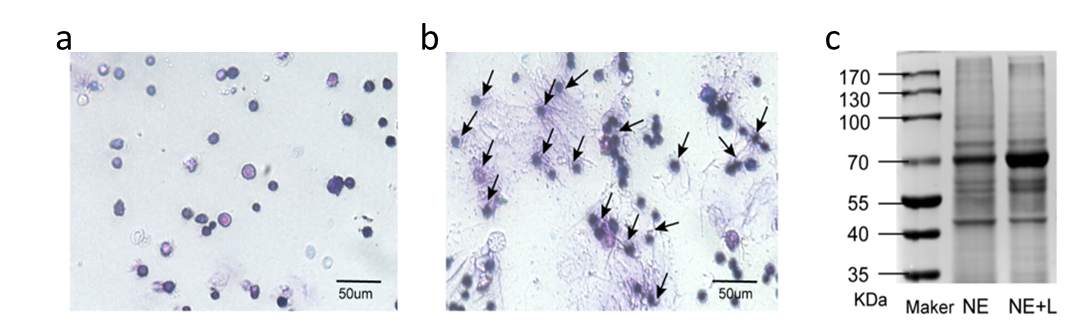

**Supplemental. Figure. 2.** The characterization of neutrophils. a. The representative image of Giemsa-stained inactivated neutrophils (NE). B. The representative image of Giemsa stained LPS activated neutrophils (NE+L). c. SDS-PAGE protein identification photograph of NE and NE+L.

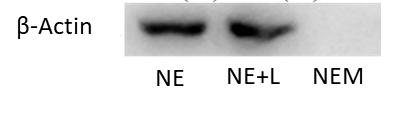

**Supplemental. Figure. 3.** The representative proteins of cytoskeleton (β-actin) in neutrophil (NE), LPS treated neutrophil (NE+L) and isolated neutrophil membrane (NEM) .

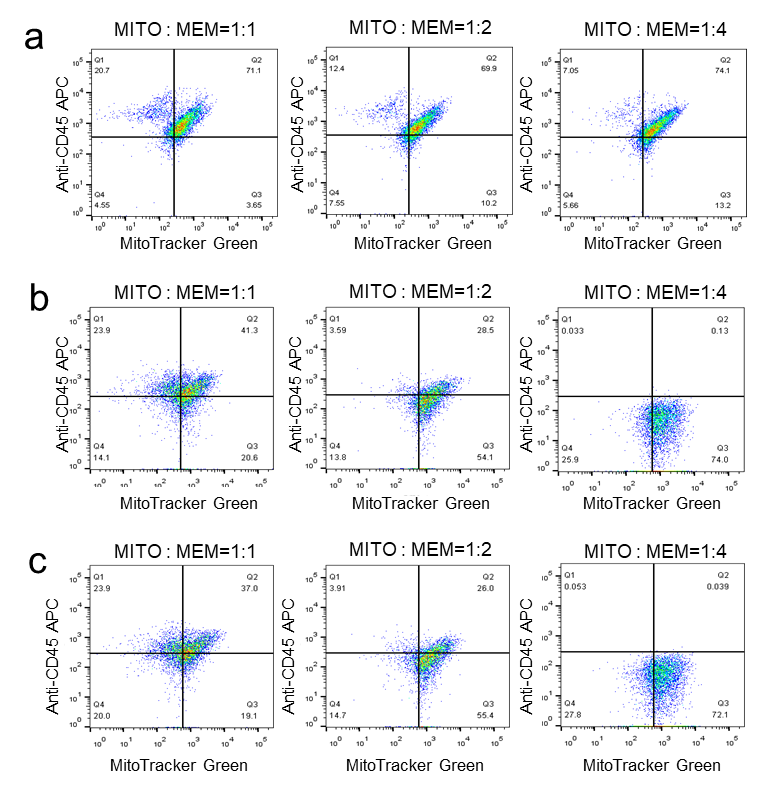

**Supplemental. Figure. 4.** The screening of the preparation method of engineered mitochondria. a. ice bathing with ultrasound for 2 min. b. Incubating on ice for 15 min. c. Incubating on ice for 30 min.

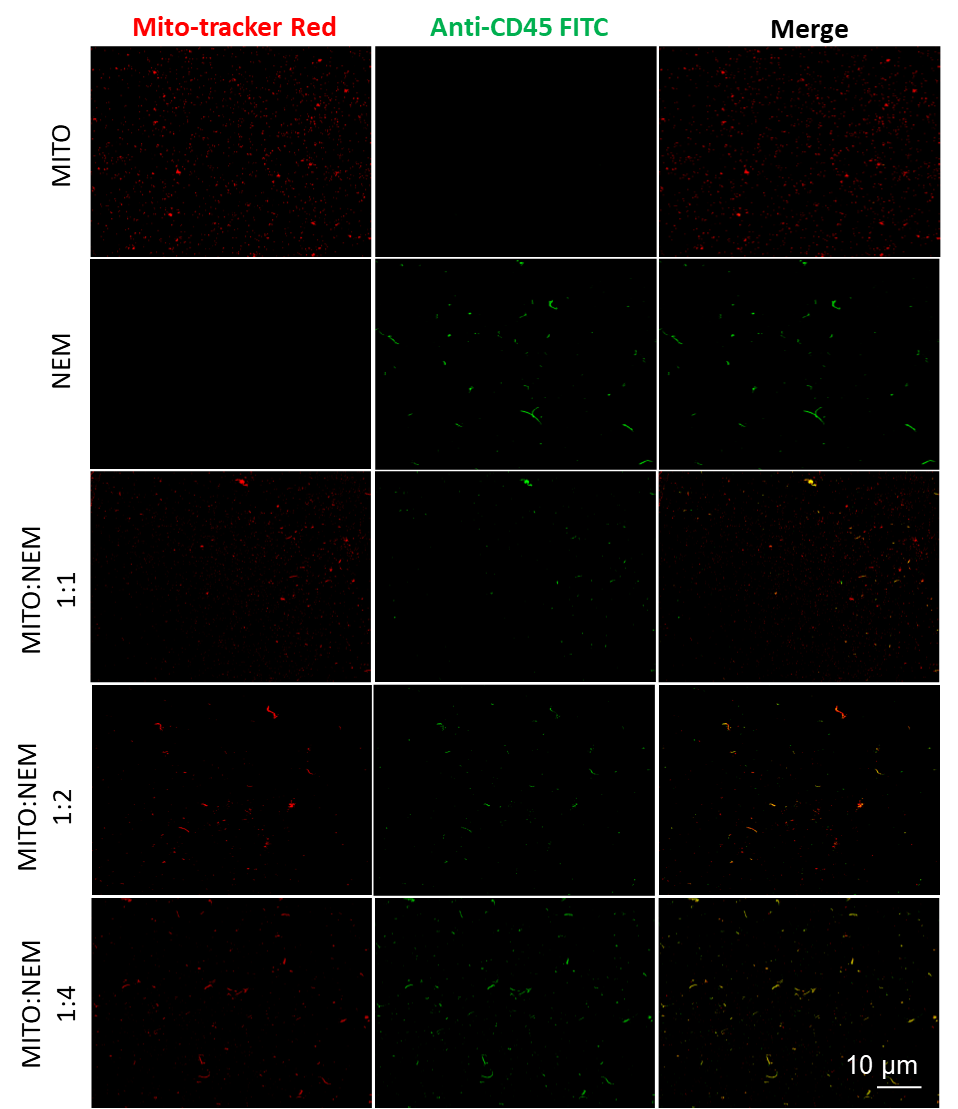

**Supplemental. Figure. 5.** The representative images of CD45 expression on MITO, NEM and nMITO acquired from different MITO:NEM ratios via ice bathing with ultrasound for 2 min.

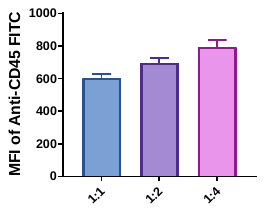

**Supplemental. Figure. 6.** The CD45 expression on nMITO acquired from different MITO:NEM ratios via ice bathing with ultrasound for 2 min determined by FACS.

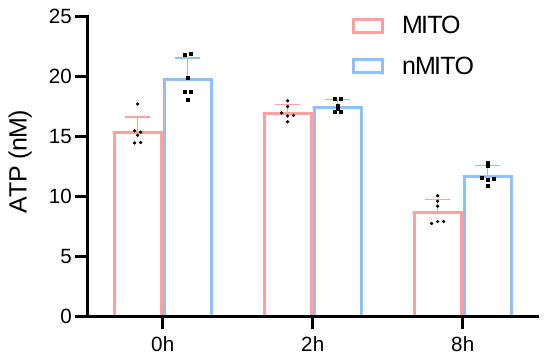

**Supplemental. Figure. 7.** Attenuation of ATP synthesis ability of mitochondria with time.

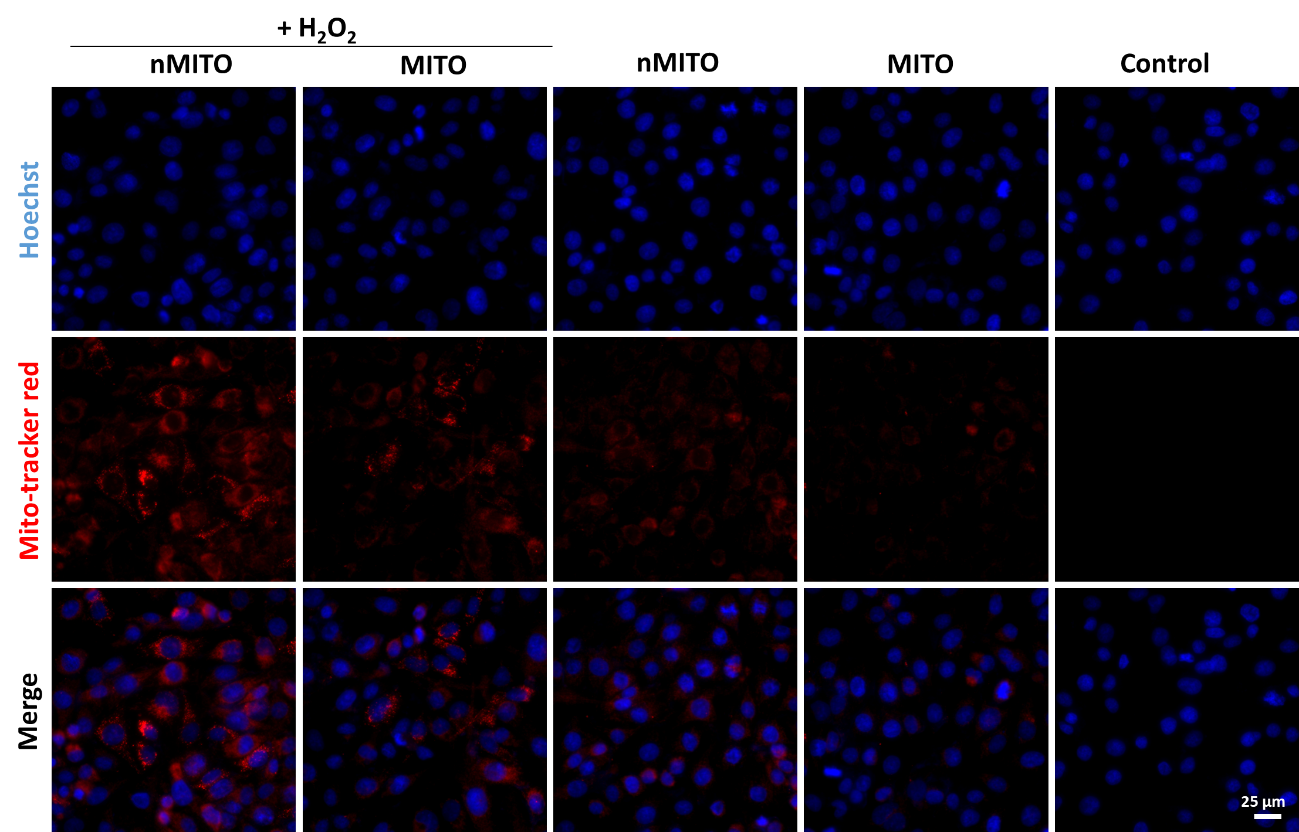

**Supplemental. Figure. 8.** The uptake of nMTIO or MITO by HUVECs cells determined by CLSM. Scale bar means 25 μm.

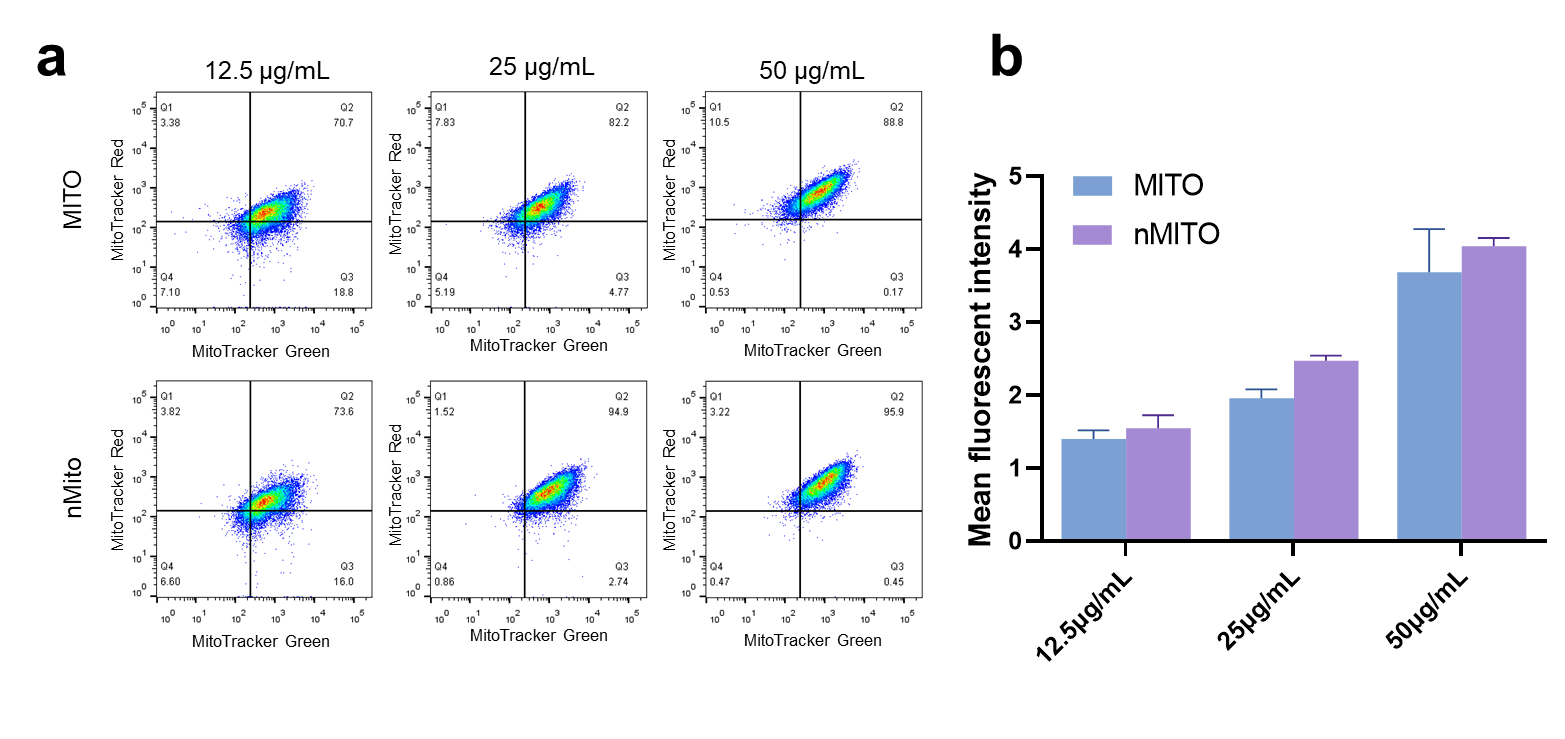

**Supplemental. Figure. 9.** The uptake of nMITO or MITO by injured HUVEC cells at different concentration determined by FACS. a. The representative fluorescence distribution of MitoTracker^®^ Red (Comp-PE-A channel, indicating the number of mitochondria) and DiO Comp-B515-A channel, indicating DiO labeled HUVEC cells). b. The mean fluorescent intensity of MITO-Tracker after injured HUVECs received nMITO or MITO.

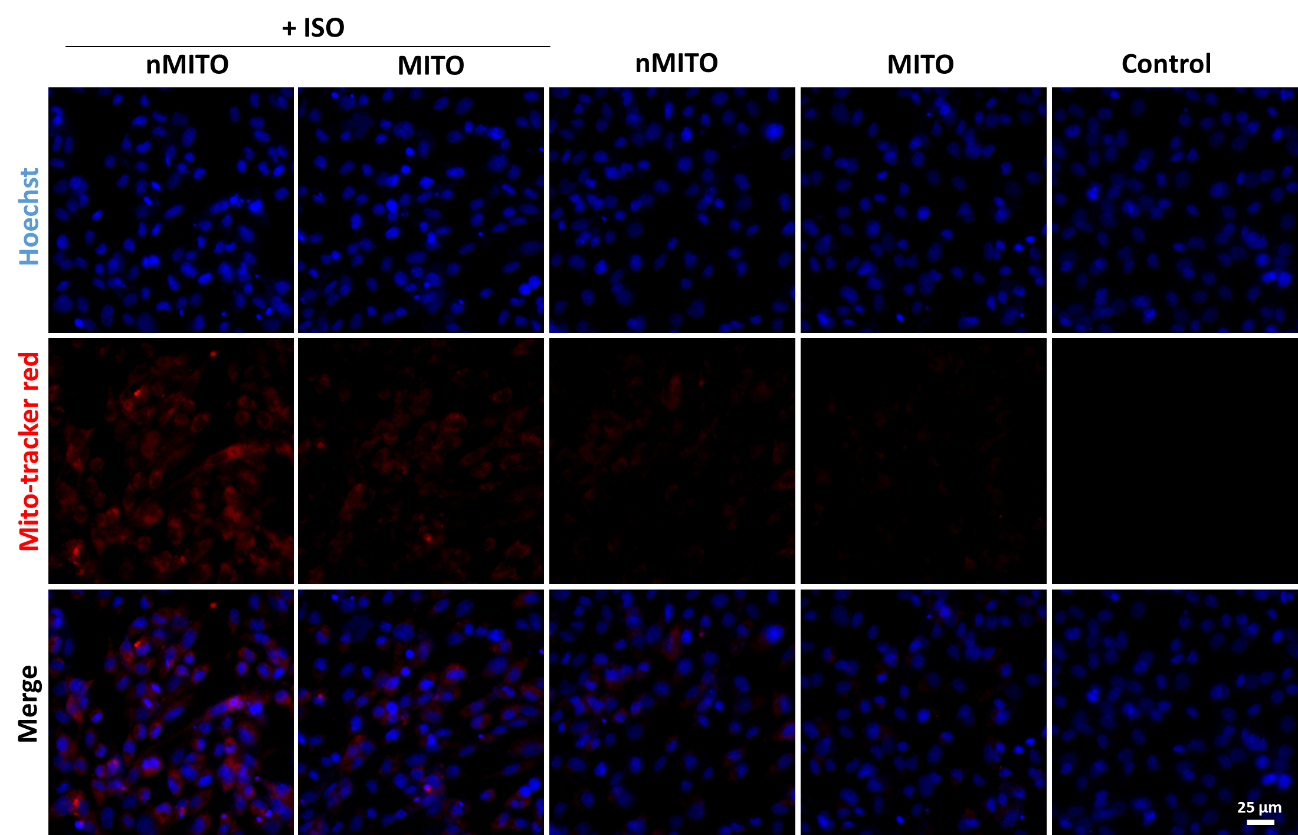

**Supplemental. Figure. 10.** The uptake of nMTIO or MITO by H9c2 cells determined by CLSM. Scale bar means 25 μm.

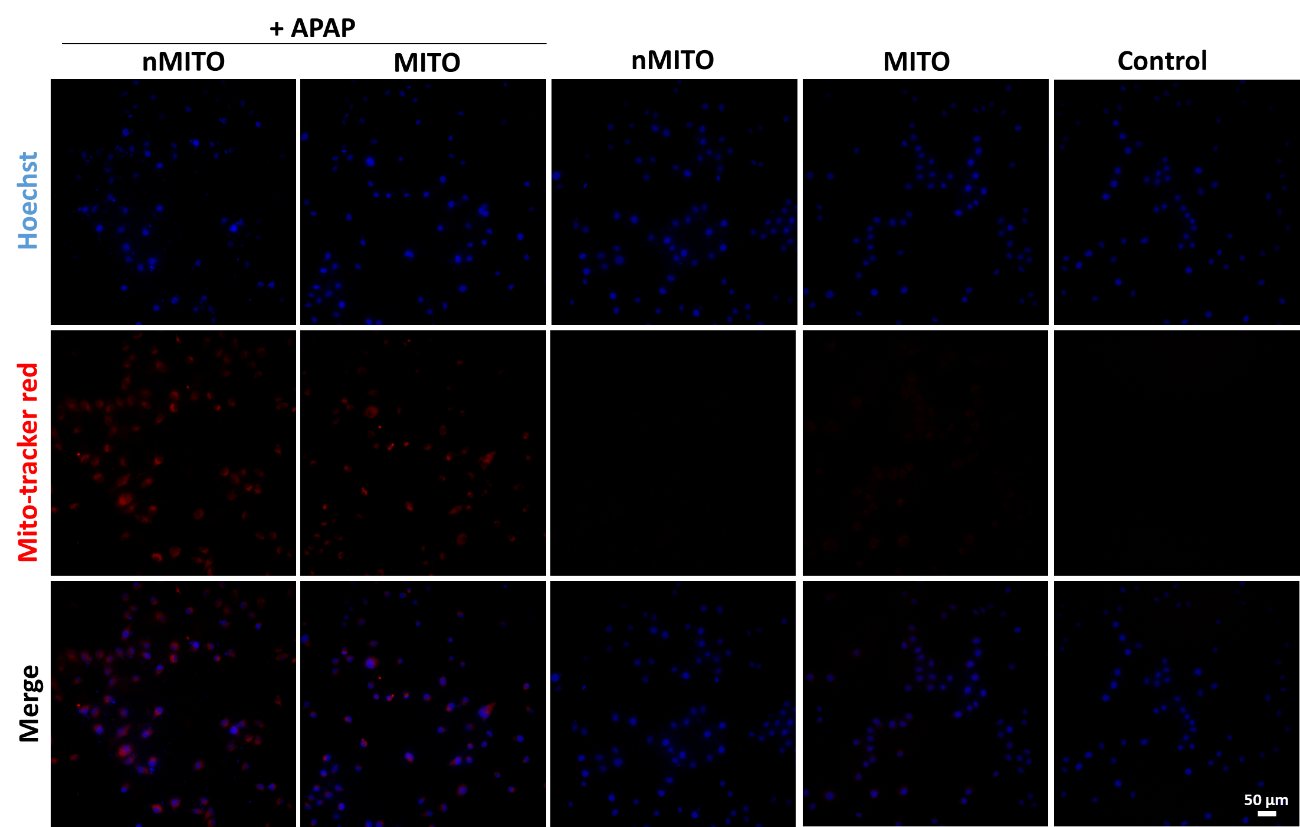

**Supplemental. Figure. 11.** The uptake of nMTIO or MITO by L02 cells determined by CLSM. Scale bar means 50 μm.

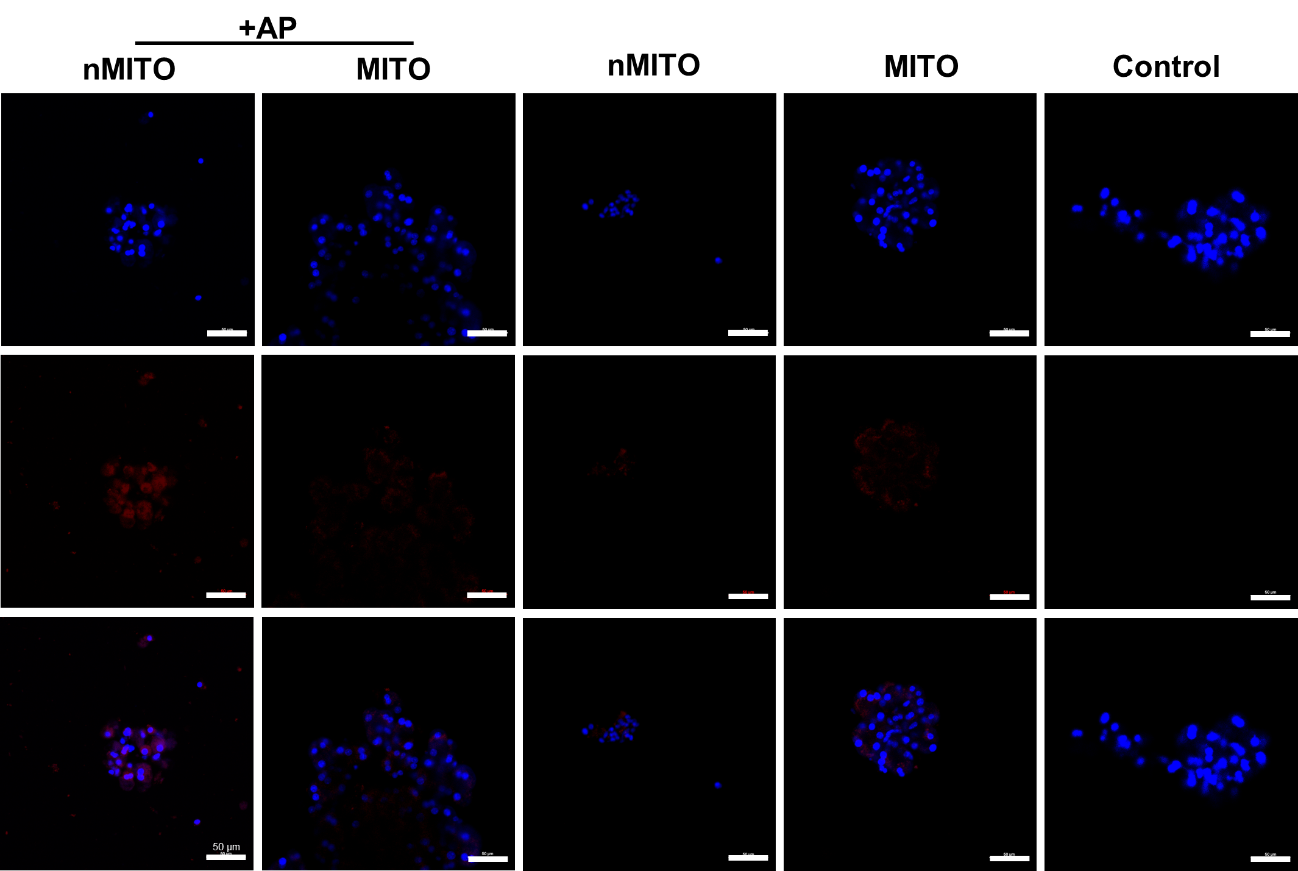

**Supplemental. Figure. 12.** The uptake of nMTIO or MITO by 266-6 cells determined by CLSM. Scale bar means 50 μm.

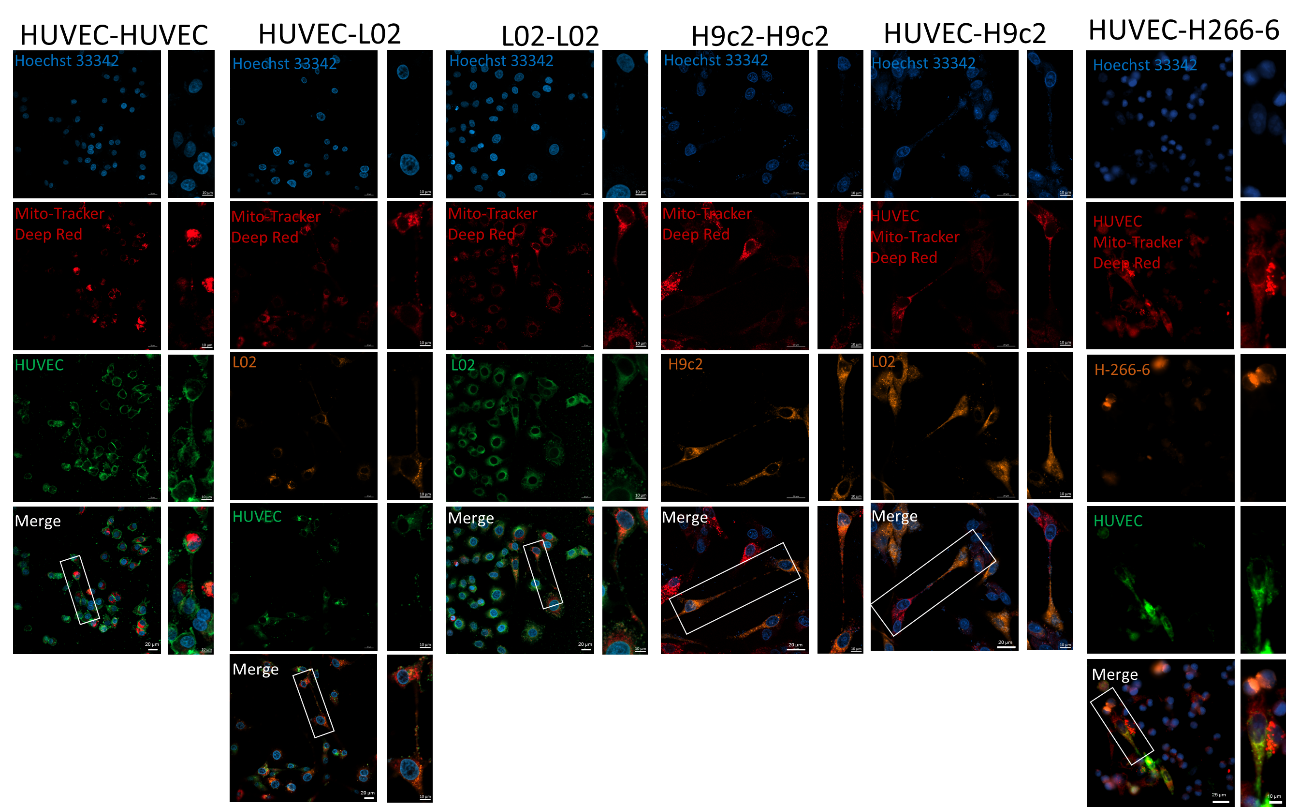

**Supplemental. Figure. 13.** The delivery of MITO form donor cell to receptor cell observed by CLSM. The scale bars in original image and partially enlarged area means 20 μm and 10 μm, respectively.

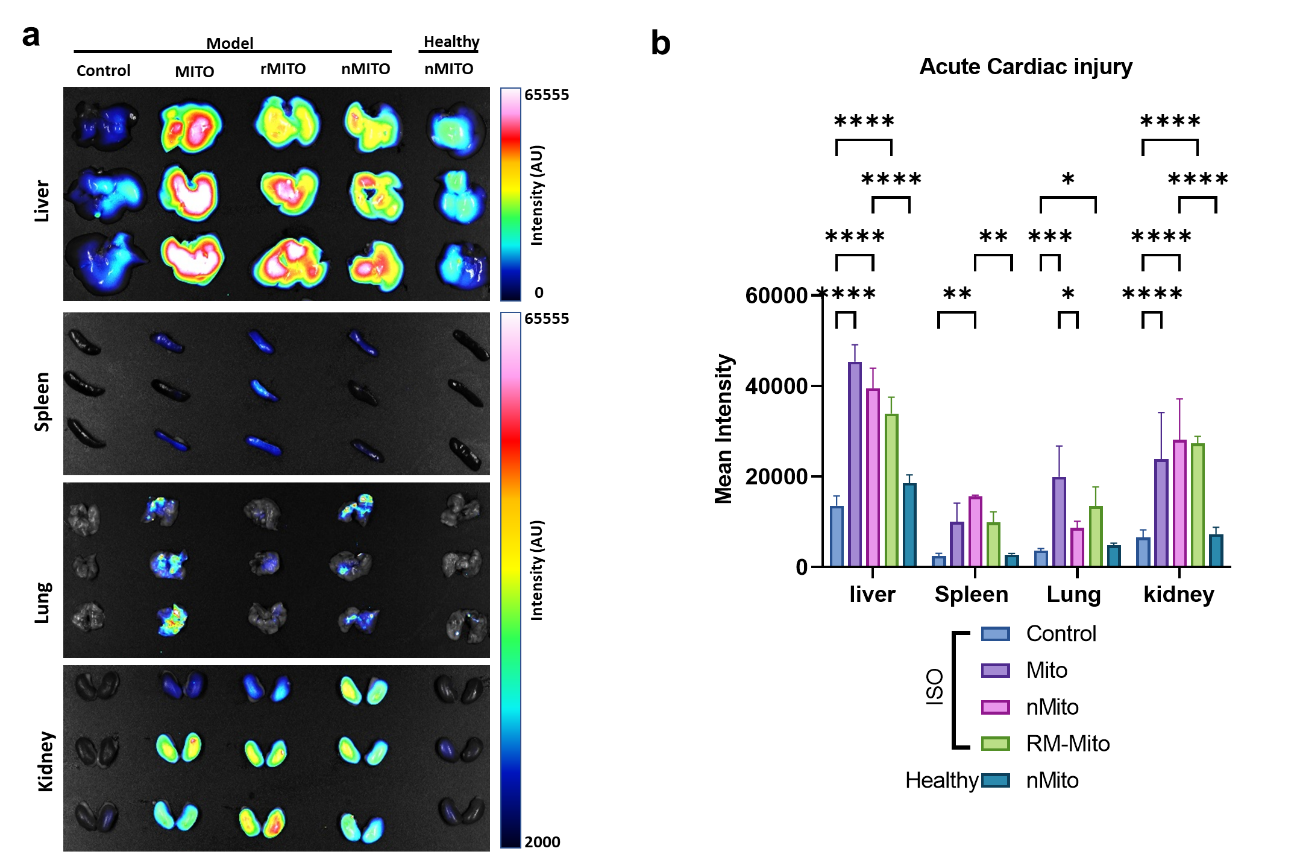

**Supplemental. Figure. 14.** The bio-distribution of MitoTracker^®^ Deep Red labeled mitochondrial in mouse with ISO induced acute cardiac injury. a. The *ex vivo* fluorescent images of main organs. b. The quantitative result of fluorescence in main organs.

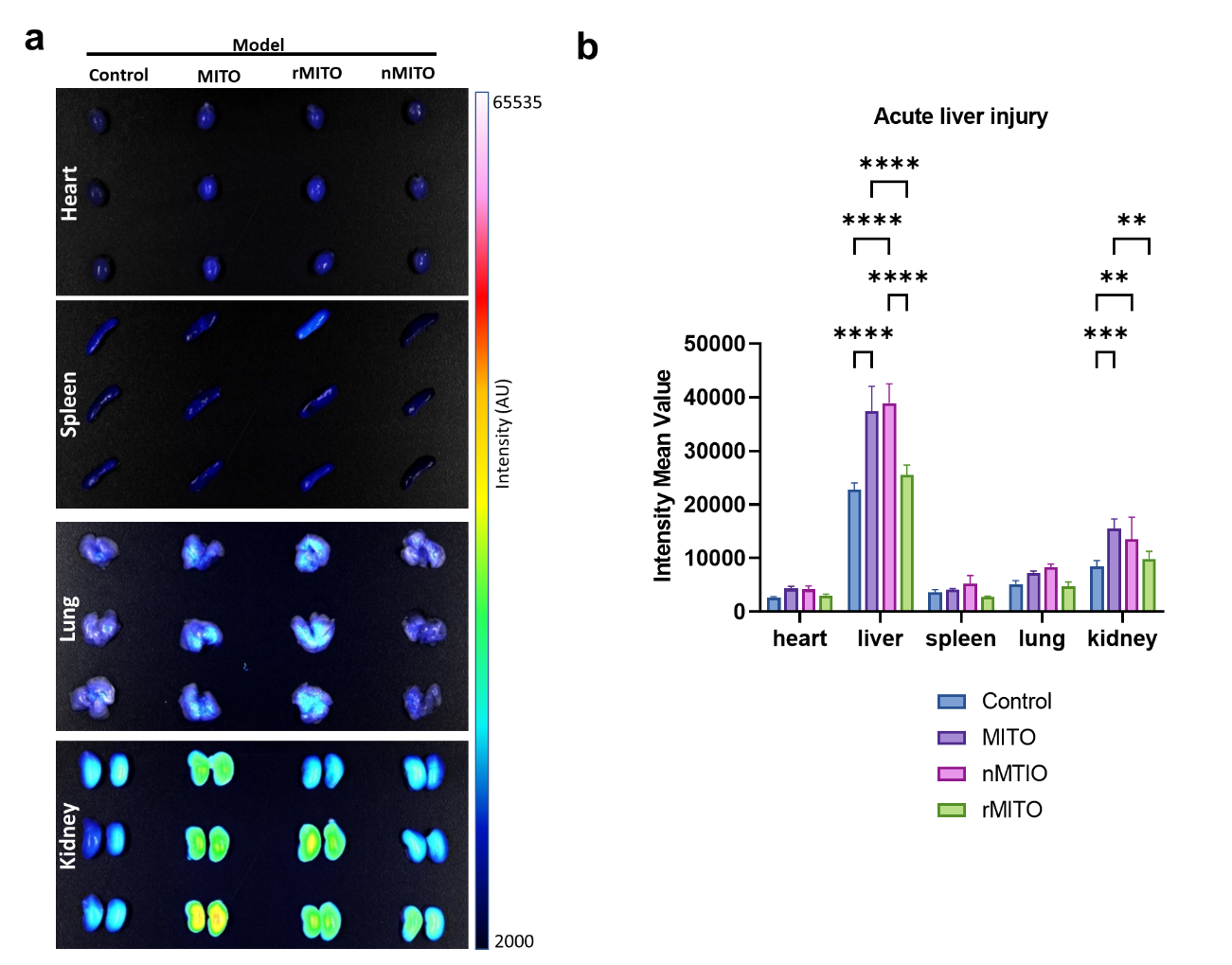

**Supplemental. Figure. 15.** The bio-distribution of MitoTracker^®^ Deep Red labeled mitochondria in mouse with APAP induced acute liver injury. a. The *ex vivo* fluorescent images of main organs. b. The quantitative result of fluorescence in main organs.

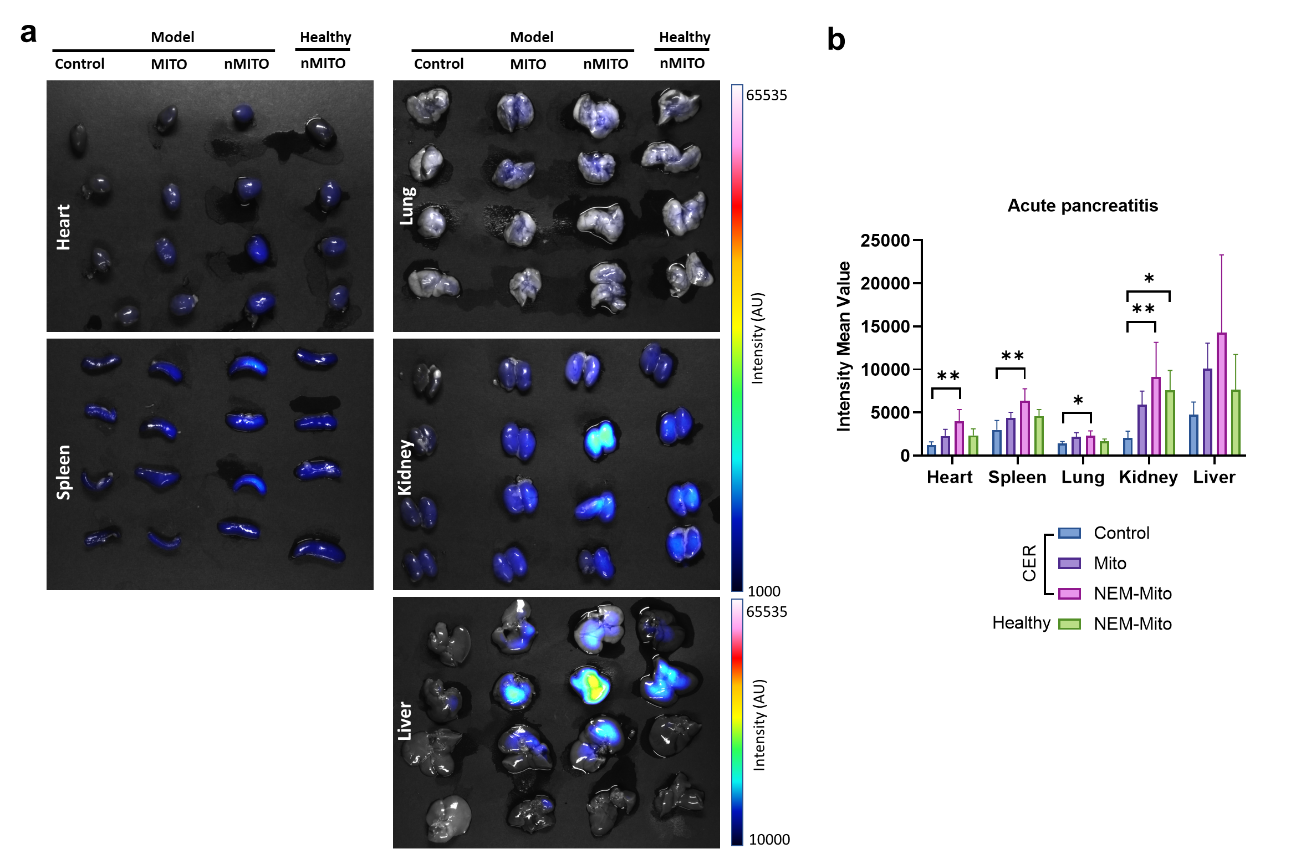

**Supplemental. Figure. 16.** The bio-distribution of MitoTracker^®^ Deep Red labeled mitochondrial in mouse with CER induced acute pancreatitis. a. The *ex vivo* fluorescent images of main organs. b. The quantitative result of fluorescence in main organs.

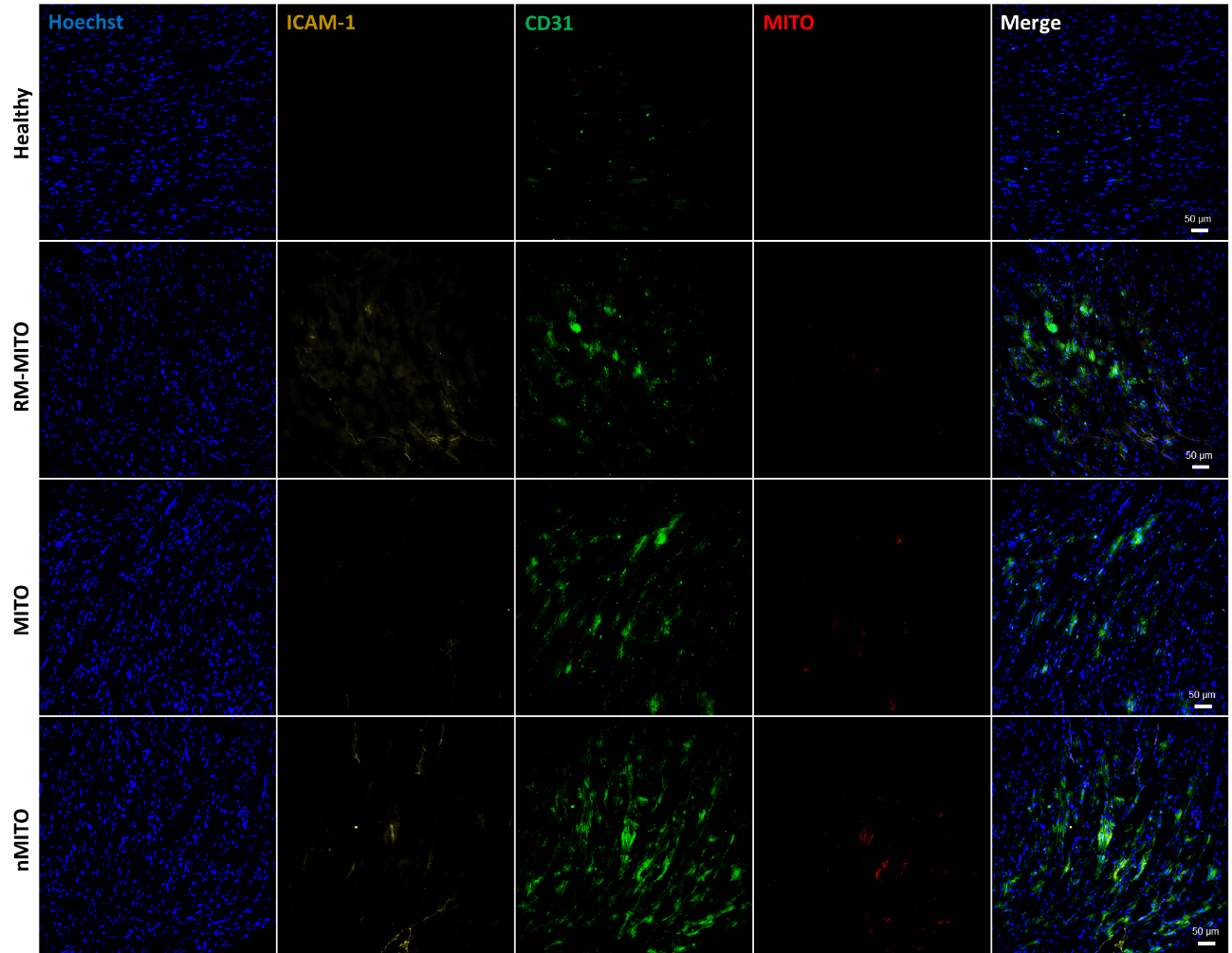

**Supplemental. Figure. 17.** Immunofluorescence images of myocardial tissue from mice bearing ISO induced acute cardiac injury. Scale bar means 50 μm.

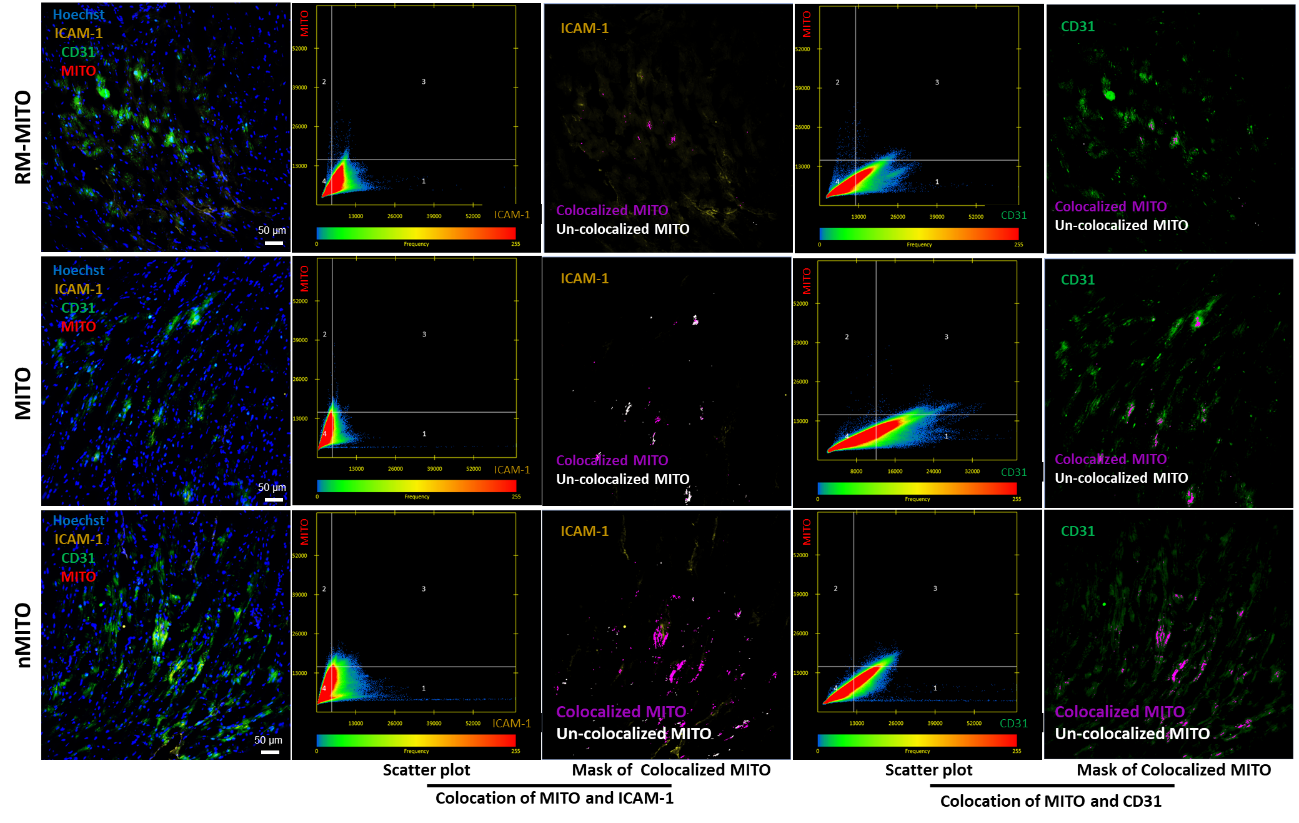

**Supplemental. Figure. 18.** The co-location analysis of mitochondria and ICAM-1/CD21 in myocardial tissue from mice bearing ISO induced acute cardiac injury. Scale bar means 50 μm.

**.**

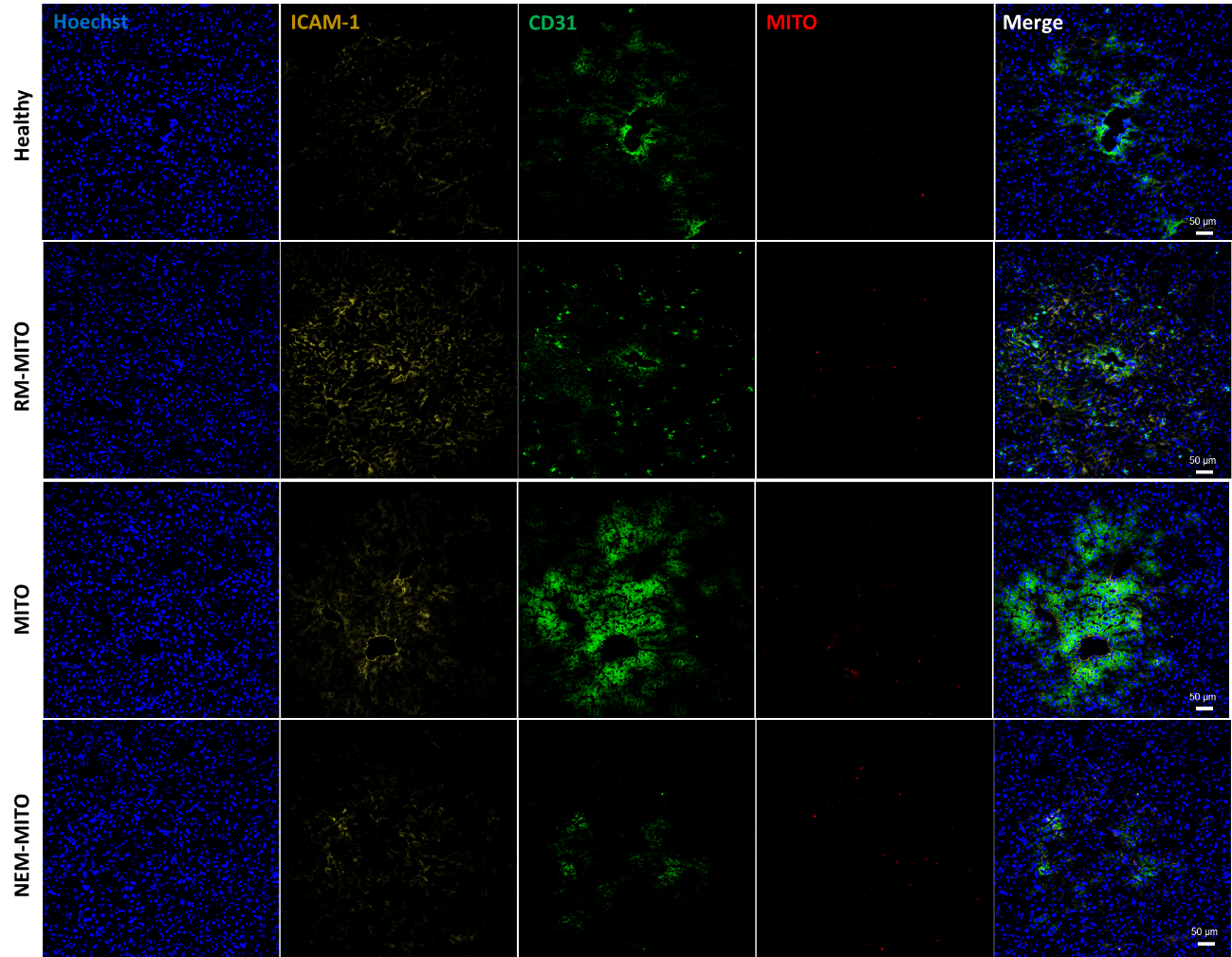

**Supplemental. Figure. 19.** Immunofluorescence images of liver tissue from mice bearing APAP induced acute liver injury. Scale bar means 50 μm.

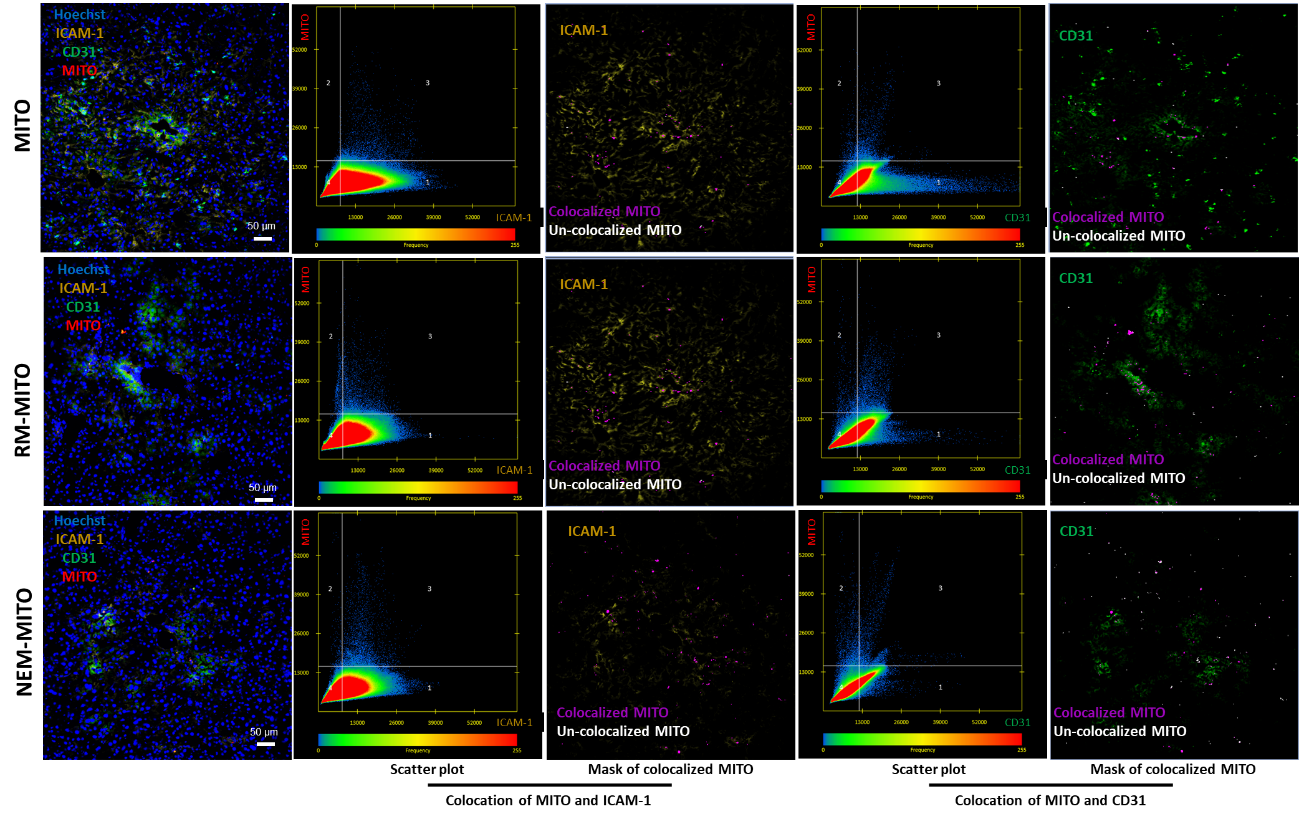

**Supplemental. Figure. 20.** The co-location analysis of mitochondria and ICAM-1/CD21 in liver tissue from mice bearing APAP induced acute liver injury. Scale bar means 50 μm.

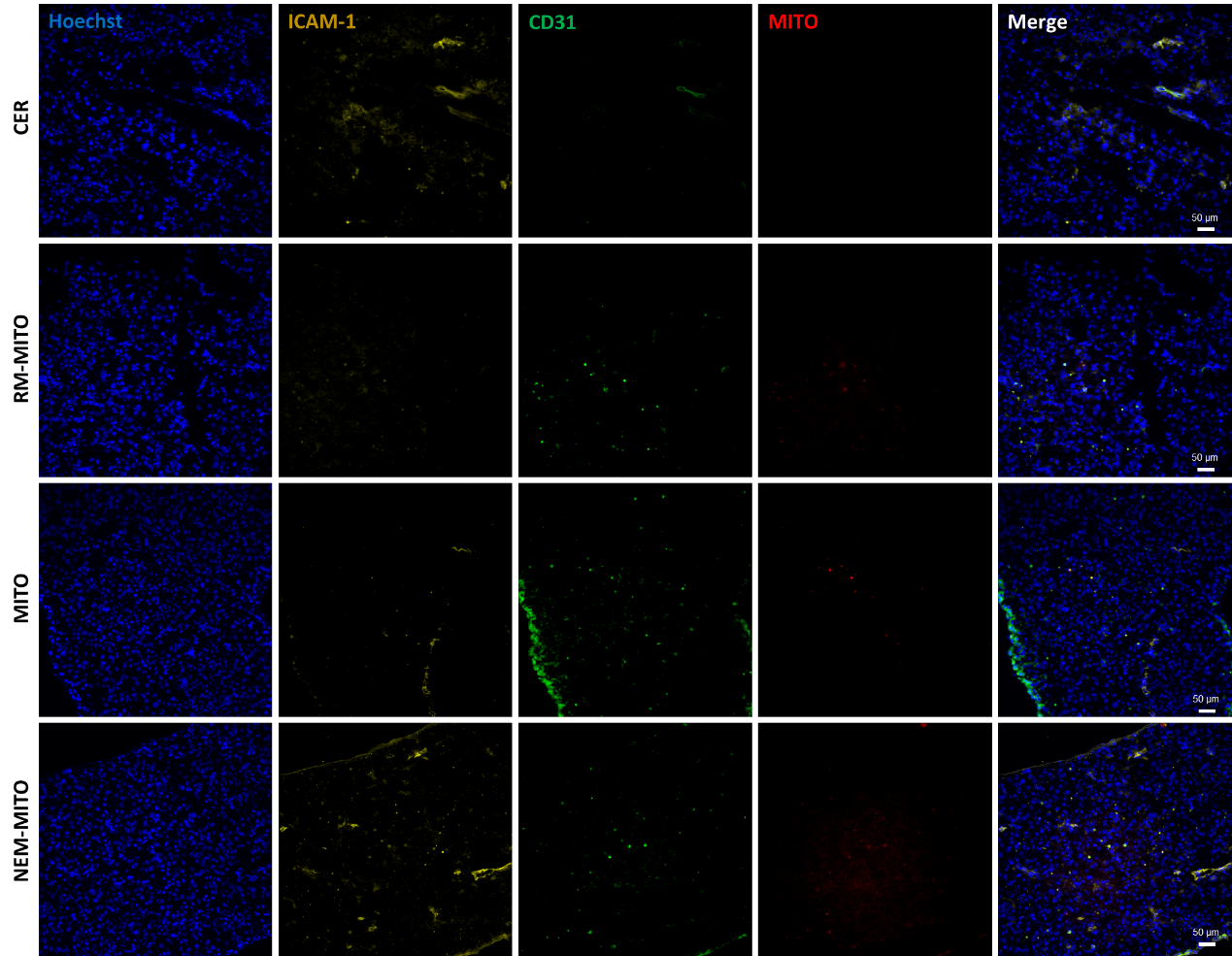

**Supplemental. Figure. 21.** Immunofluorescence images of pancreas tissue from mice bearing CER induced acute pancreatitis. Scale bar means 50 μm.

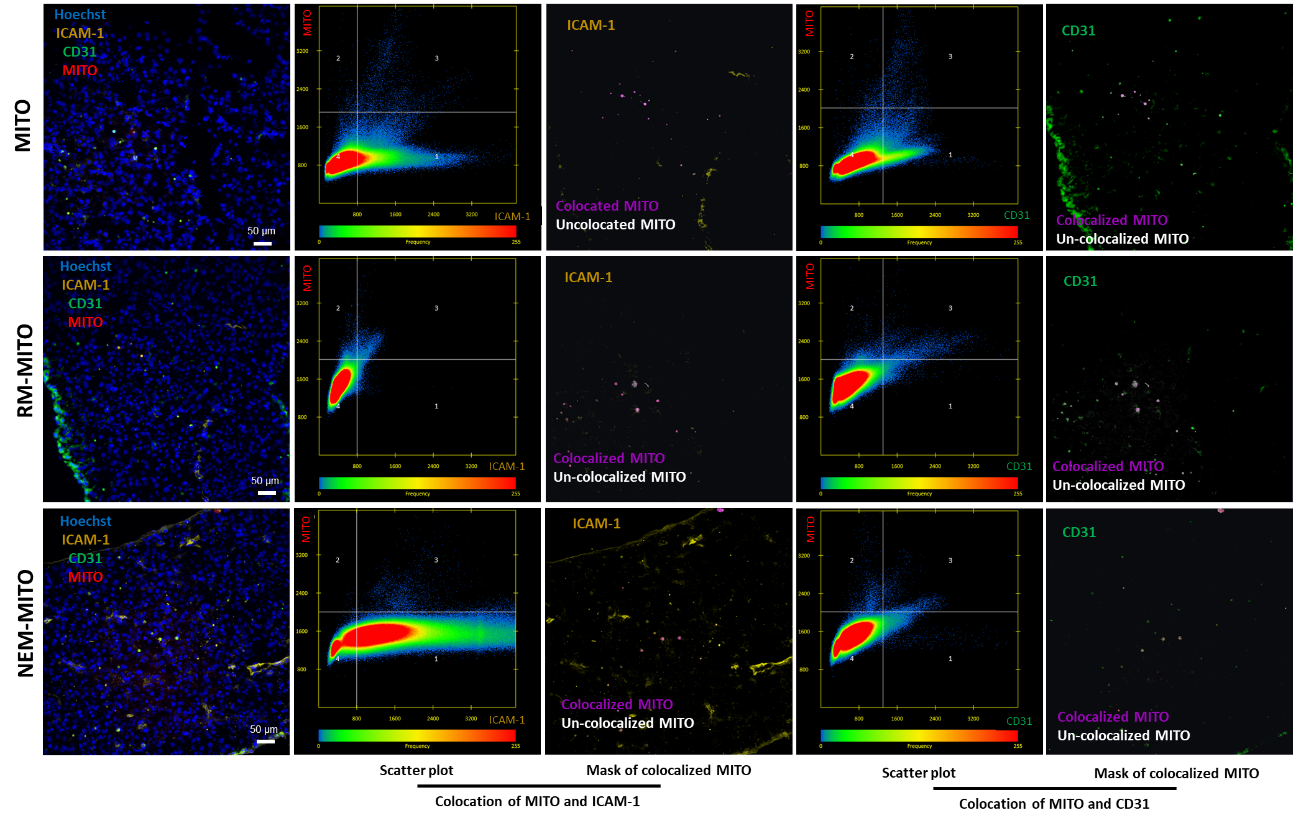

**Supplemental. Figure. 22.** The co-location analysis of mitochondria and ICAM-1/CD21 in pancreas tissue from mice bearing CER induced acute pancreatitis. Scale bar means 50 μm.

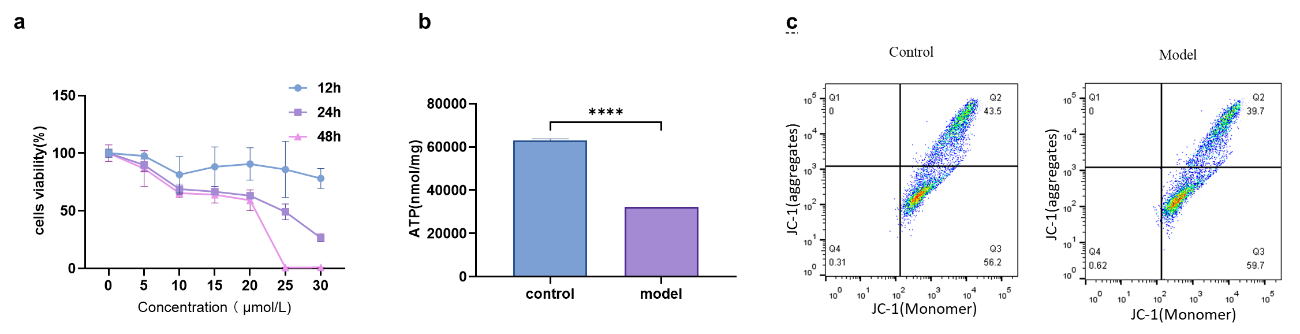

**Supplemental. Figure. 23.** The construction of isoproterenol mediated H9c2 cell injury cells. a. The cell viability of H9c2 cell under different concentration of ISO. b. The ATP synthesis level of healthy H9c2 cell (control) and injured H9c2 cell(model). c. The mitochondrial membrane potential of healthy H9c2 cell (control) and injured H9c2 cell(model).

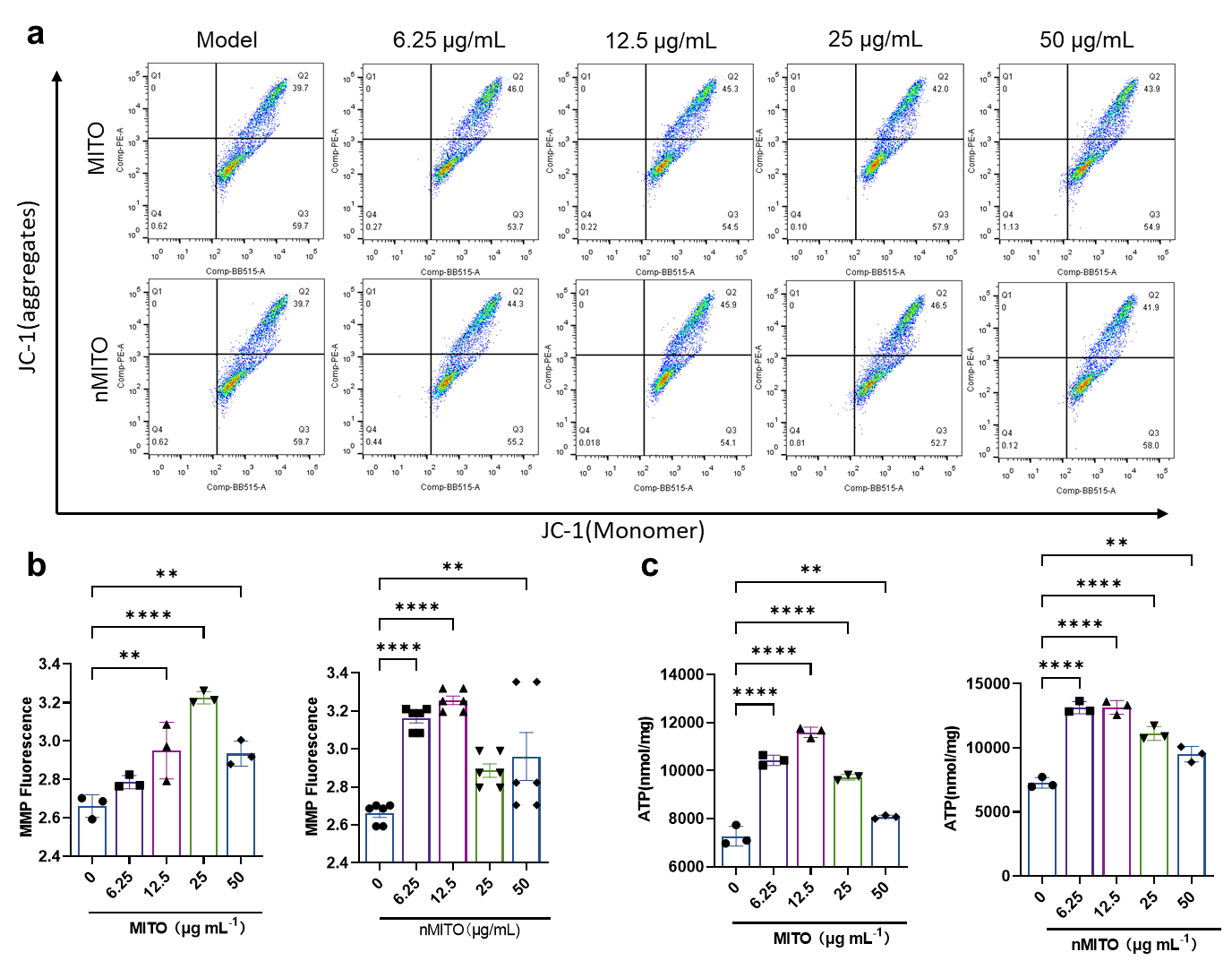

**Supplemental. Figure. 24.** The *in vitro* cell repairing activity against injured H9c2 cells of nMITO. a. Measurement of JC-fluorescence by flow cytometry, in H9c2 cells received different concentration of nMTIO or MITO. b. The mitochondrial membrane potential of H9c2 cells received different concentration of nMTIO or MITO, calculated by JC-1 Red/Green signal intensity. c. The ATP synthesis levels of H9c2 cells received different concentration of nMTIO or MITO.

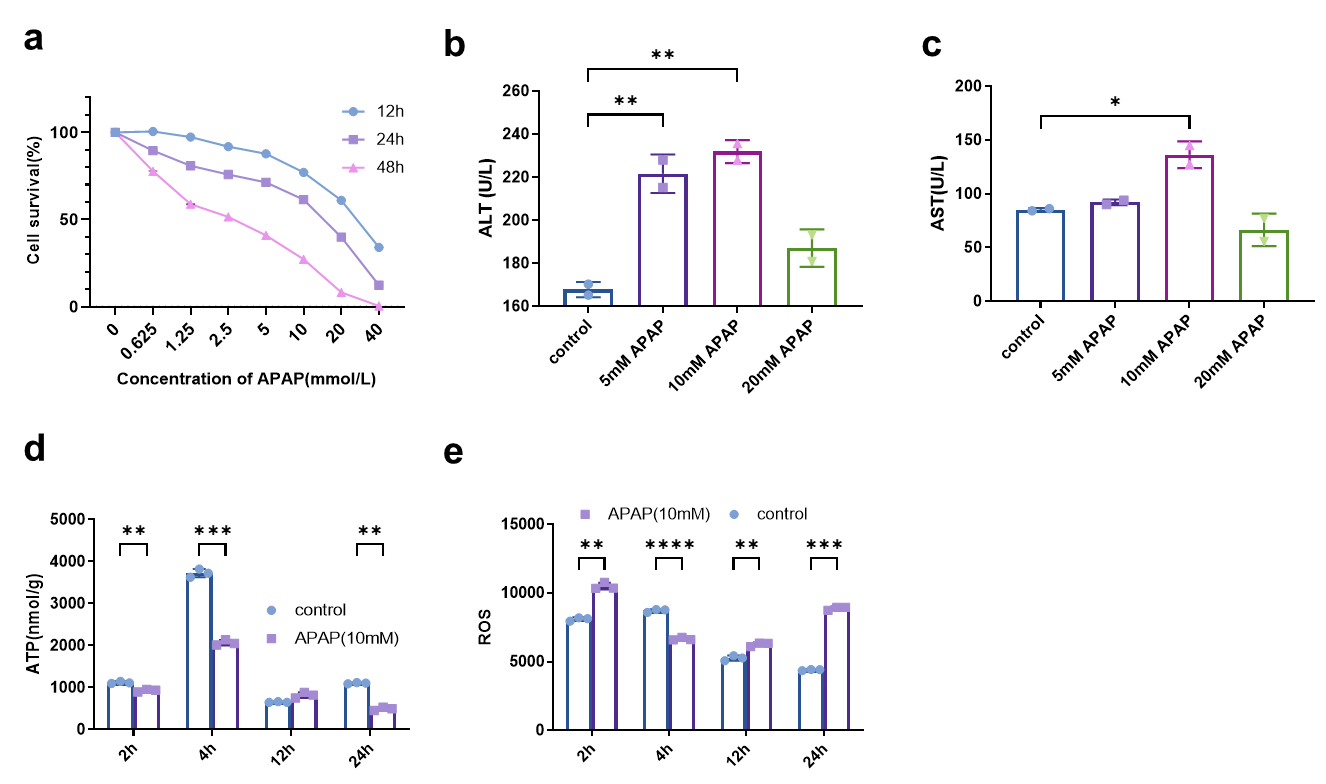

**Supplemental. Figure. 25.** The construction of L02 hepatocyte cells injured by acetaminophen (APAP). a. The cell viability of L02 cell under different concentration of APAP. b, c. The ALT(b) and AST(c) levels of L02 cell under different concentration of APAP. d,e. The ATP (d) and ROS (e) levels of injured L02 cell after stimulated by APAP with different hours.

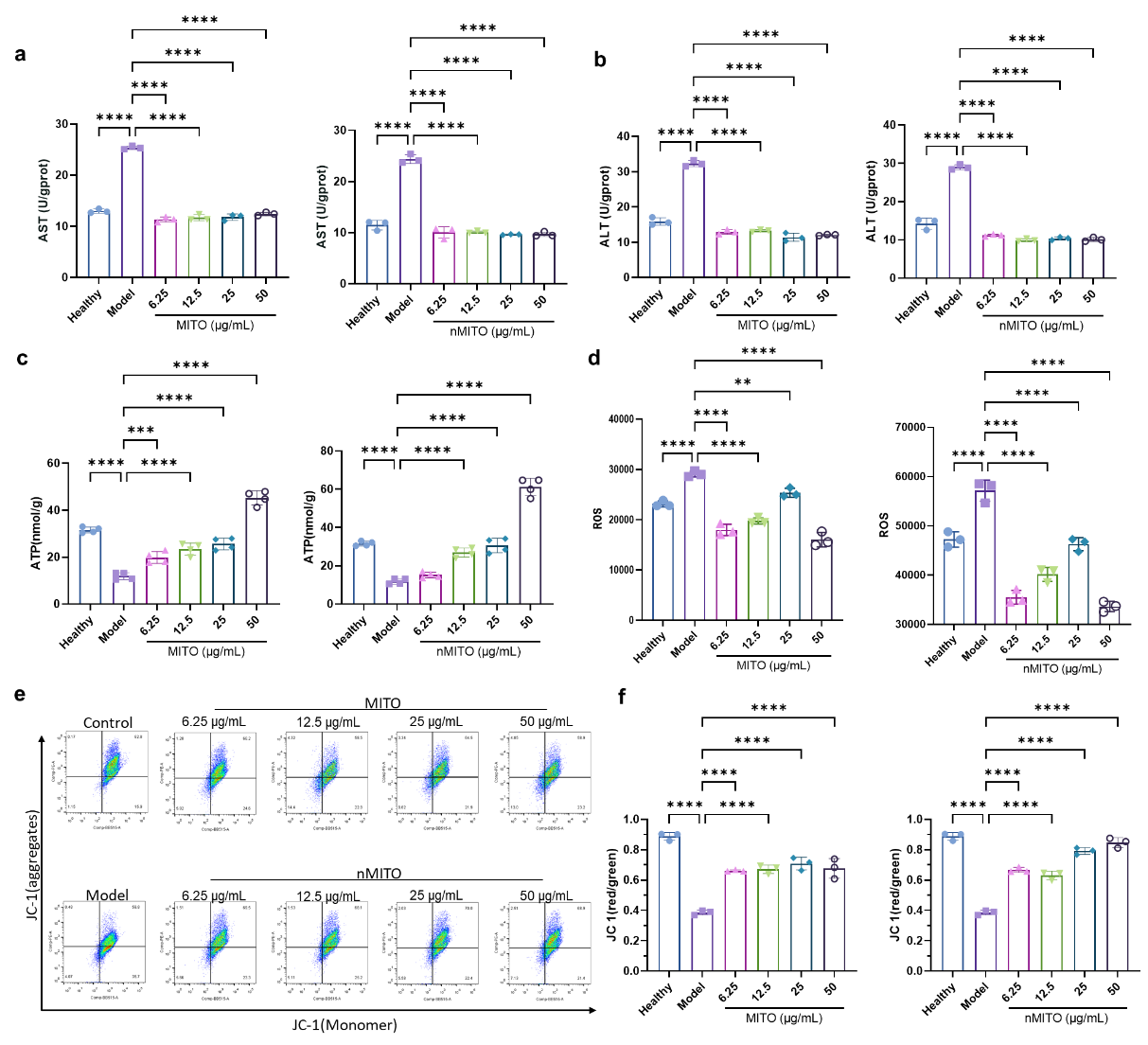

**Supplemental. Figure. 26.** The *in vitro* cell repairing activity against injured L02 cells of nMITO.

**a-d.** The ALT(a), AST(b), ATP synthesis (c), and ROS(d) levels of APAP-injured-L02 cells received different concentration of MITO or nMTIO. e. Measurement of JC-fluorescence by flow cytometry, in L02 cells received different concentration of MITO or nMTIO. f. The mitochondrial membrane potential of L02 cells received different concentration of MITO or nMTIO, calculated by JC-1 Red/Green signal intensity.

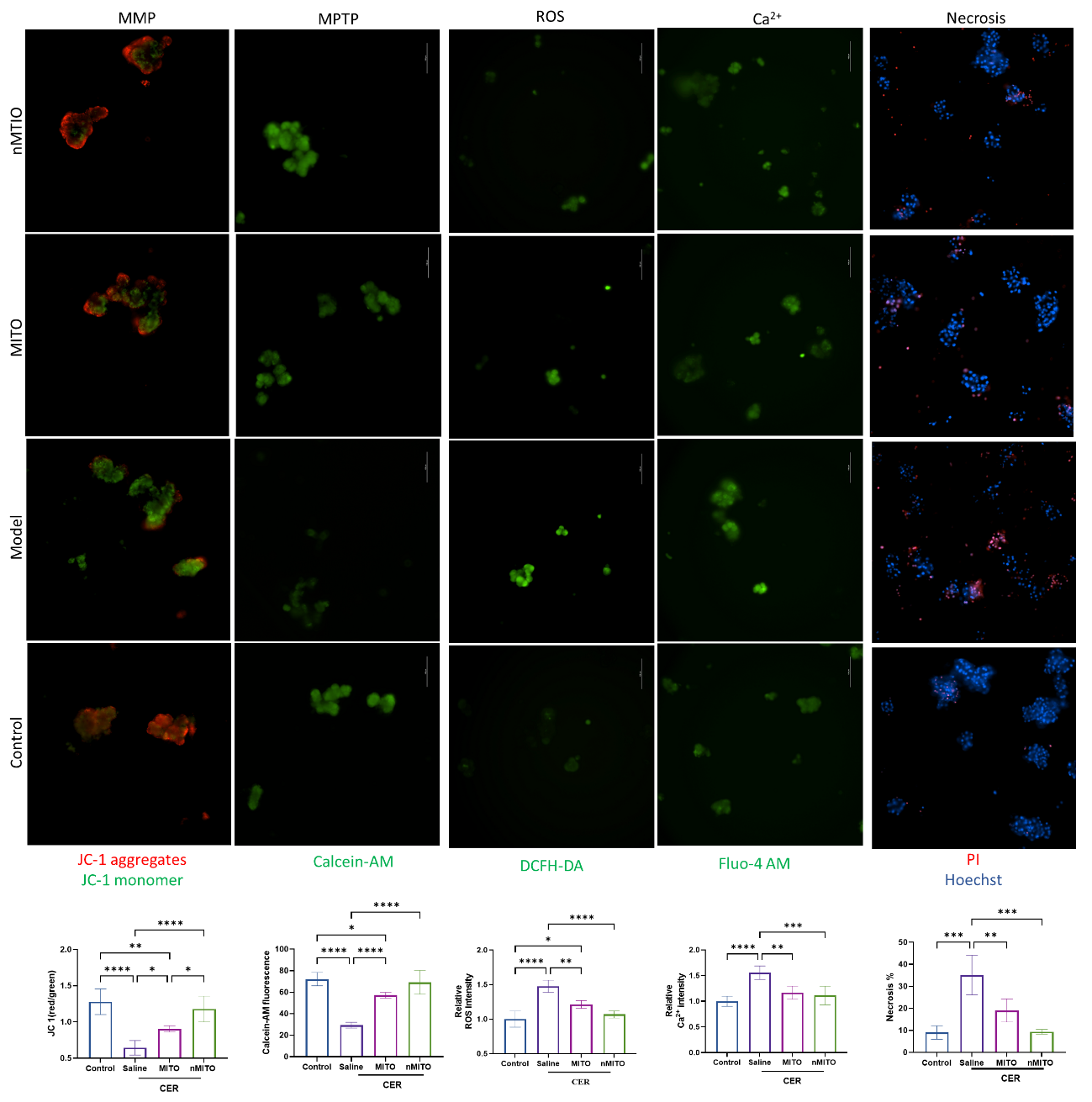

**Supplemental. Figure. 27.** The determination and quantitative results of mitochondrial membrane potential (MMP), mitochondrial permeability transition pore (MPTP) opening, ROS, relative Ca^2+^ intensity and necrotic rate of CER treated acinar cells after receive saline or nMTIO.

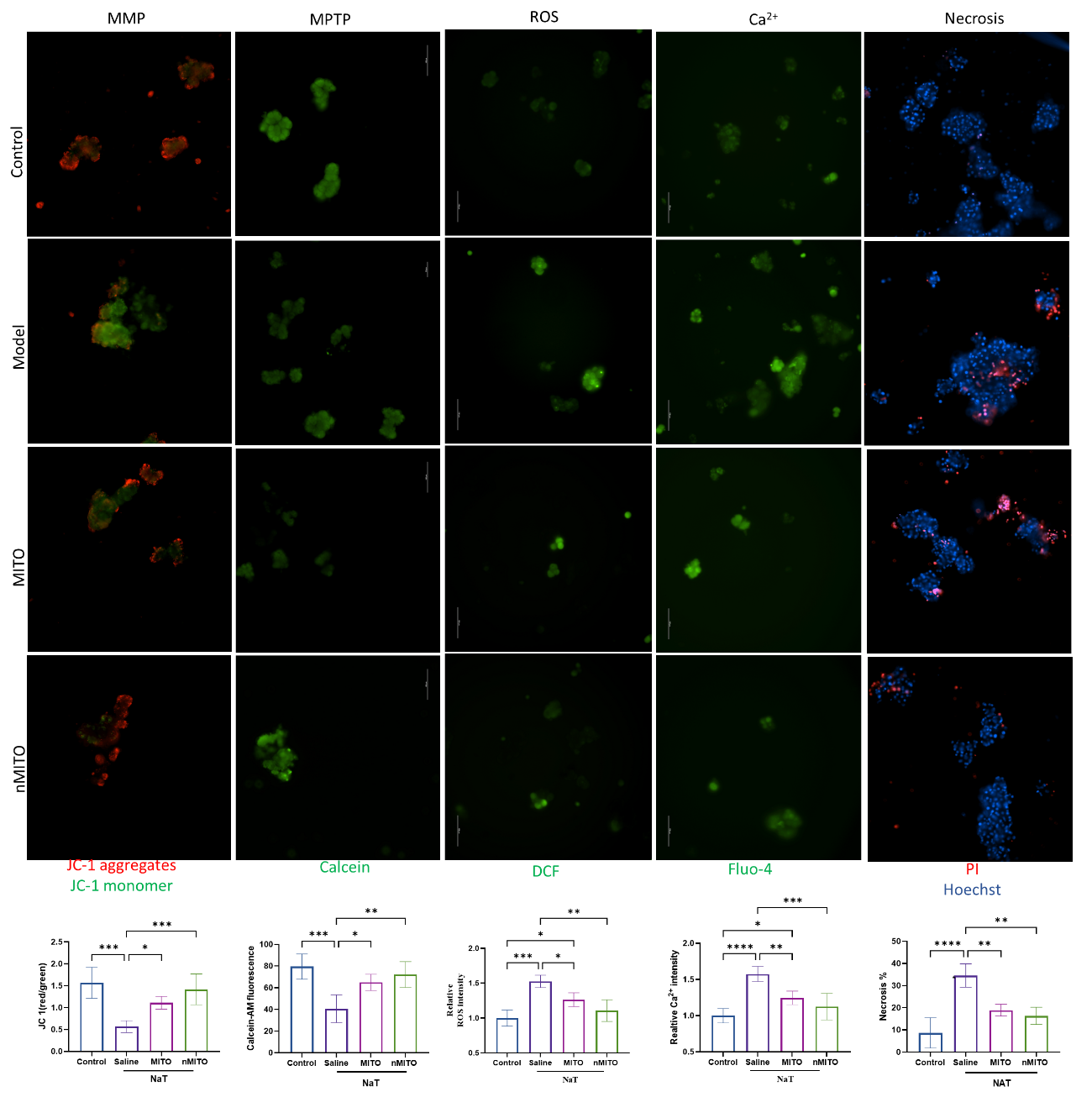

**Supplemental. Figure. 28.** The determination and quantitative results of mitochondrial membrane potential (MMP), mitochondrial permeability transition pore (MPTP) opening, ROS, relative Ca^2+^ intensity and necrotic rate of CER treated acinar cells after receive saline or nMTIO.

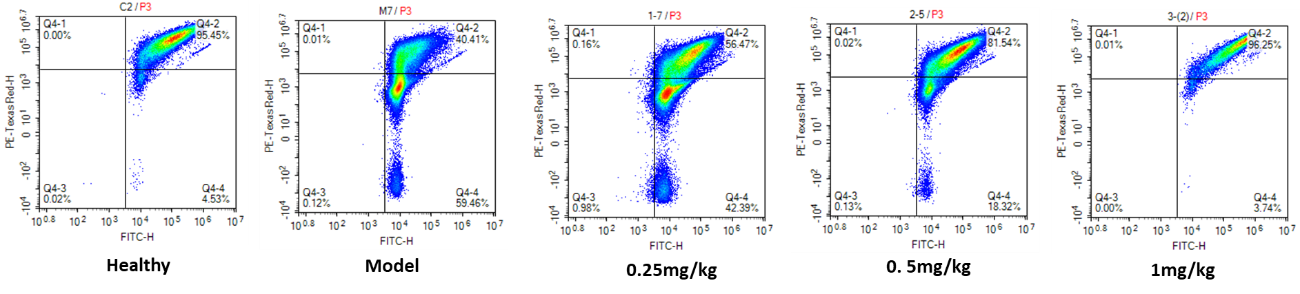

**Supplemental. Figure. S29.** The mitochondrial membrane potential of mouse cardiomyocytes in heart tissue of mice after treatment.

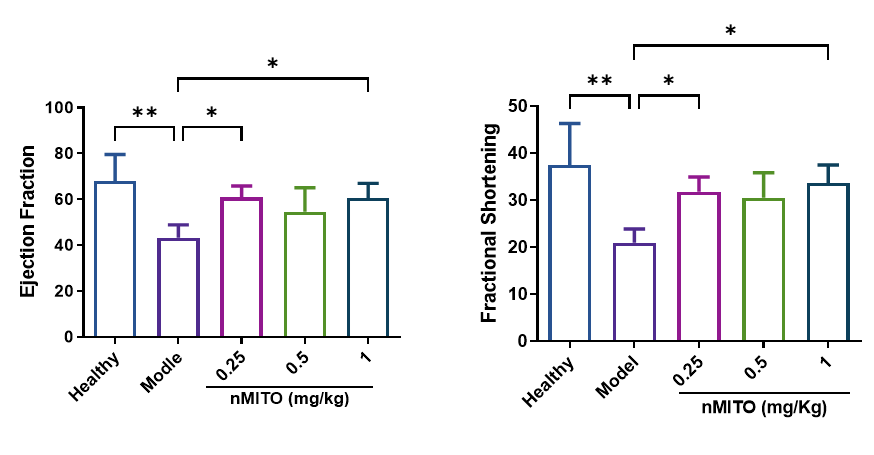

**Supplemental. Figure. 30.** The ejection fraction (EF) and fractional shortening (FS) of mice treated with nMITO at dose of 0.25 or 1 mg/Kg).

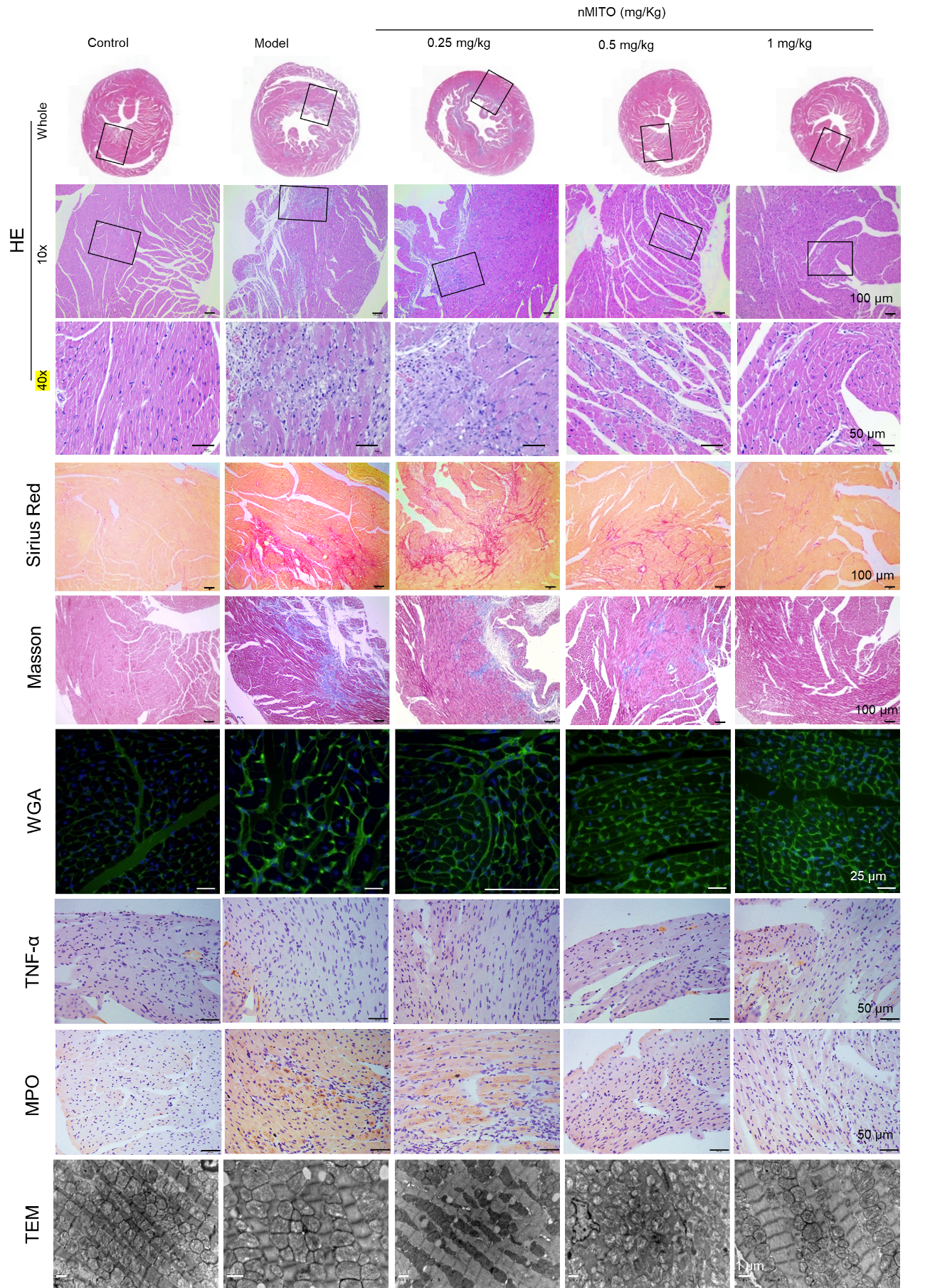

**Supplemental. Figure. 31.** The results of hematoxylin-eosin (HE) staining, fibrosis expression, inflammatory infiltration and mitochondrial appearance in the myocardium of mice after treatment.

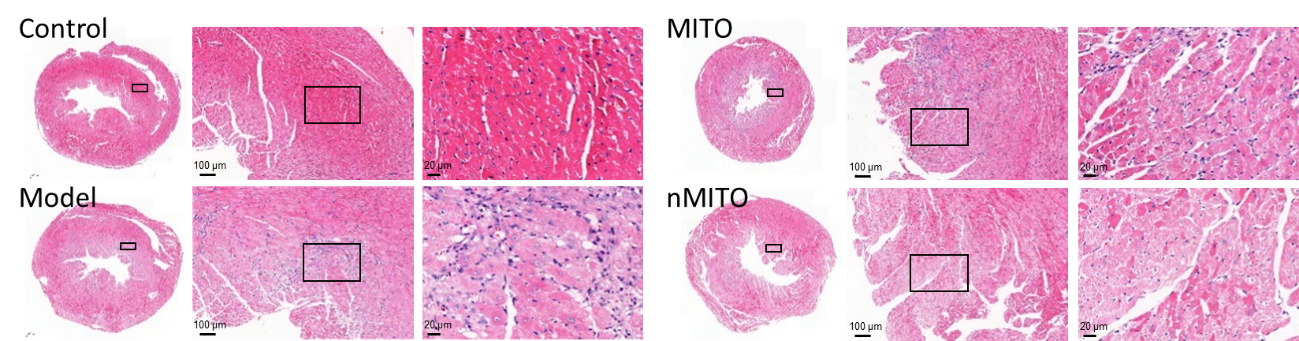

**Supplemental. Figure. 32.** The hematoxylin-eosin (HE) staining of myocardium of mice after treatment with saline, MITO or nMITO.

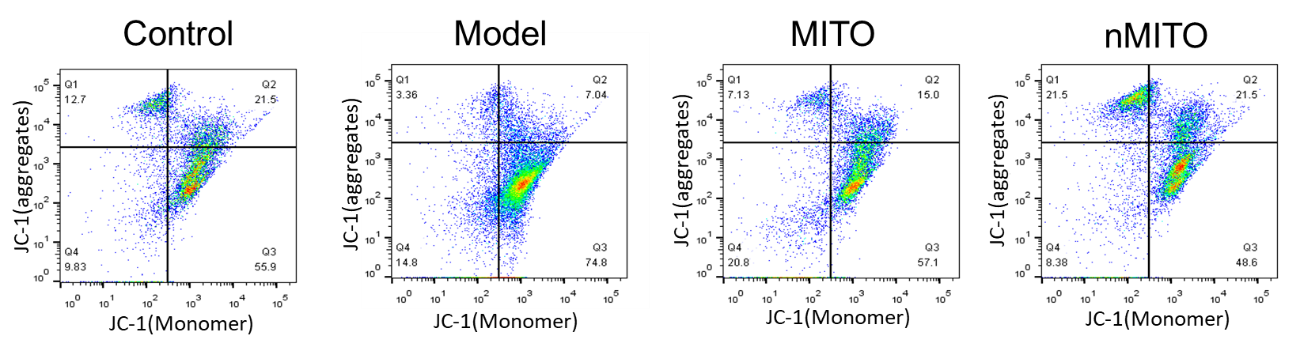

**Supplemental. Figure. 33.** The mitochondrial membrane potential of mouse cardiomyocytes in heart tissue of mice after treatment.

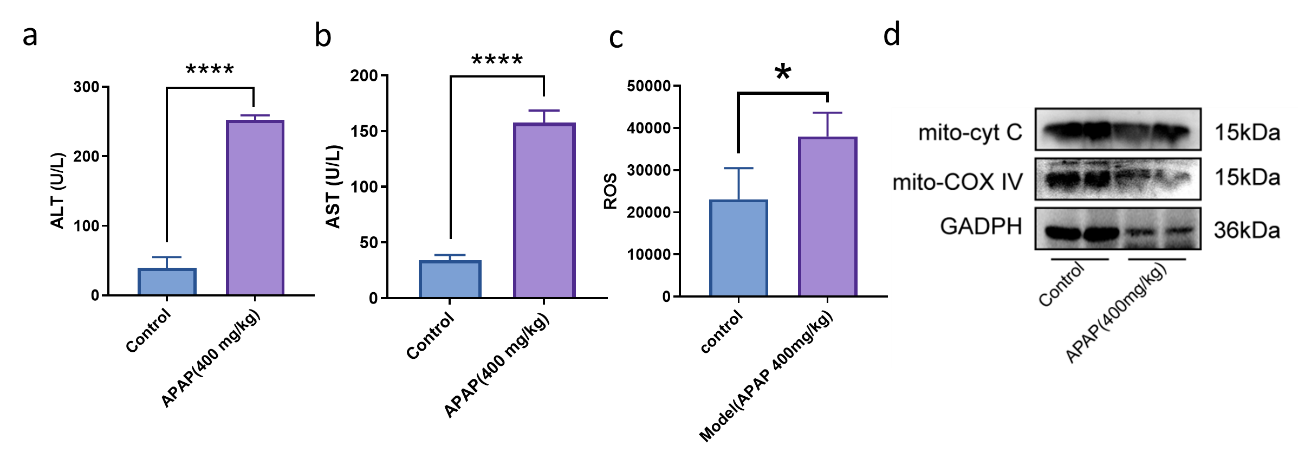

**Supplemental. Figure. 34.** The construction of acetaminophen (APAP)-mediated acute liver injury. a,b. Results of liver function indexes such as ALT (a) and AST(b).c. Evaluation of ROS activities of hepatocytes. d. The Western blot results for mitochondrial-associated proteins.

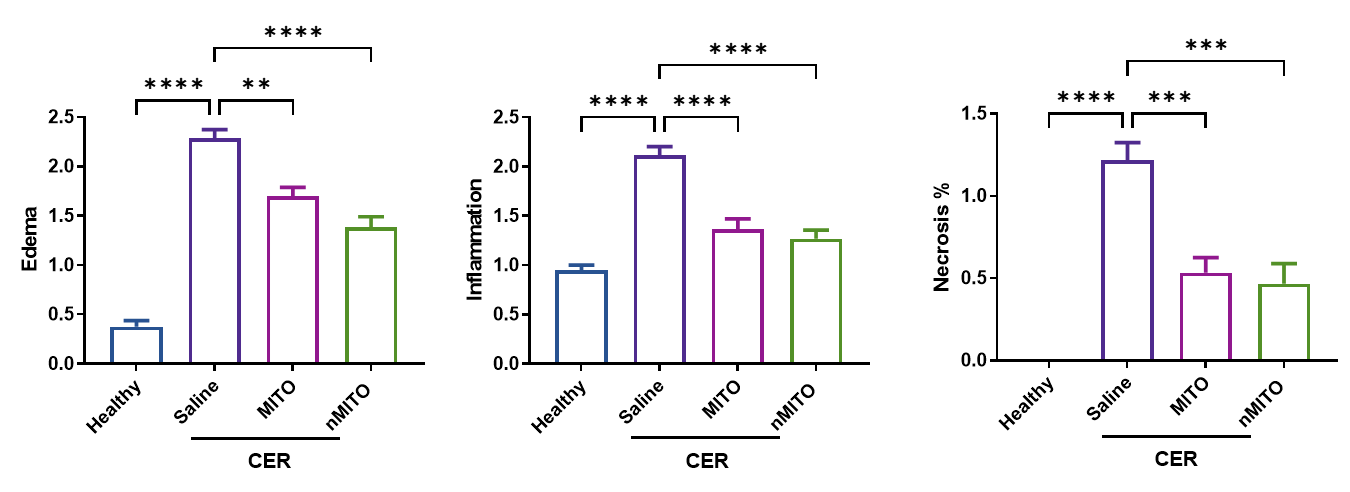

**Supplemental. Figure. 35.** Pathological scoring of edema, inflammation, and necrosis in pancreatic H&E sections from CER-AP mice.

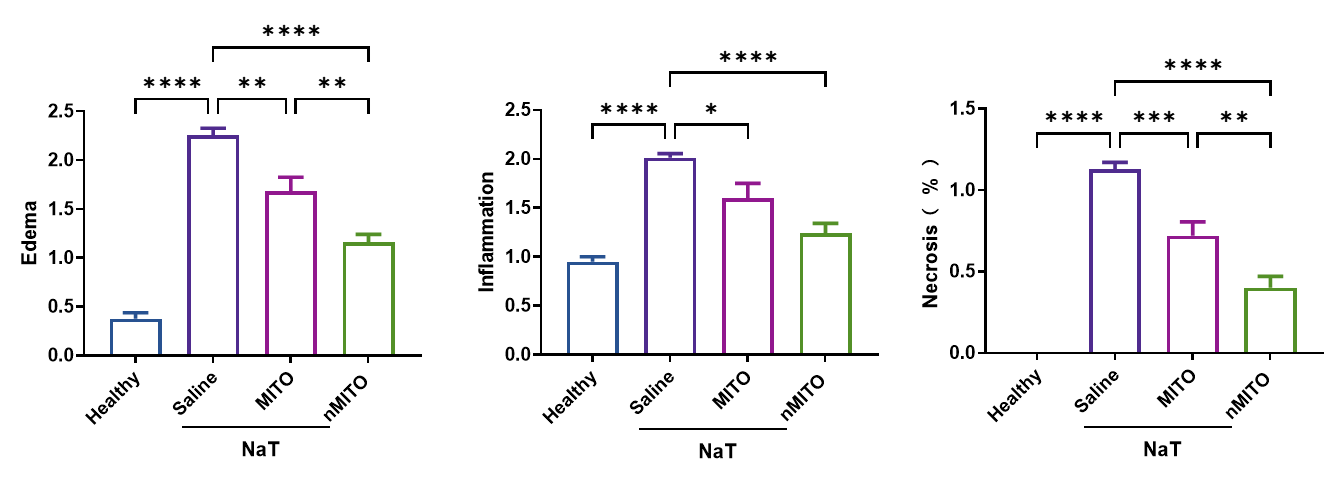

**Supplemental. Figure. 36.** Pathological scoring of edema, inflammation, and necrosis in pancreatic H&E sections from NaT-AP mice.

.
